# Maximal human lifespan in light of a mechanistic model of aging

**DOI:** 10.64898/2025.12.22.695887

**Authors:** Ben Shenhar, Shachaf Frenkl, Tomer Levy, Uri Alon

## Abstract

Why has maximal human lifespan barely changed in the past two centuries? To understand this we make a mechanistic link between cellular damage, survival curves, and maximum lifespan using a validated stochastic model of damage accumulation and extensive human data. We show that maximal lifespan is set mainly by damage production and clearance rates, as in progeroid syndromes. In contrast, lifestyle factors such as exercise, nutrition, and sleep chiefly reduce stochastic noise and raise the damage level compatible with survival, shifting the median but not the maximum. Similar constraints arise in other mortality models. Our analysis predicts that lifestyle can extend maximal lifespan by at most ∼1 year; substantial gains will require directly perturbing damage production or removal, suggesting specific molecular targets.

## Introduction

Maximal human lifespan has remained nearly unchanged, despite a doubling of median life expectancy in the past 150 years(*1–4*). For instance, life expectancy at age 30 rose by 20 years but life expectancy at age 90 only rose by 2-3 years(*5–7*). Over the past century, survival curves have generally become more rectangular, as early deaths are prevented and deaths cluster closer to the maximal lifespan^1^ but historical changes appear to have almost no effect on maximal lifespan. In contrast, maximal lifespan can be readily extended in model organisms such as in yeast, C. elegans, fruit flies, and mice, adding to the puzzle in humans.

There is a gap in understanding the rigidity of maximal human lifespan in terms of molecular and cellular mechanisms of aging. Despite advances in characterizing the molecular and cellular changes with age in humans and model organisms, it is unclear which mechanisms affect median and maximal lifespan differentially. For example, although it is often thought that aging is due to accumulating damage(*8*), it is not known how median and maximal lifespan are differentially affected by damage production rate, removal rate, stochastic noise and threshold for death. Understanding which factors affect maximal lifespan may offer clues for future longevity interventions.

To address this, we applied an advance that links mechanistic aging processes to demographic variables such as median and maximal lifespan. This advance is a mathematical model of stochastic damage accumulation called the Saturating-Removal (SR) model(*9*, *10*). The SR model was developed based on dynamics of senescent cells in mice(*10*) and has since been shown to capture a wide range of aging patterns including the exponential rise in hazard with slowdown at old age(*9*, *10*), exponential disease incidence(*11*), effect of parabiosis(*12*), longevity interventions and their combinations(*13*), and aging differences between species(*14*). Recently, the model has been used to reexamine the heritability of human lifespan(*15*) and provide insights into the compression of morbidity(*13*).

The model considers damage accumulation as a balance between production that rises with age, removal that saturates at high damage and stochastic noise. Death occurs when damage crosses a threshold. The model thus has four key parameters - production rate, removal rate, noise amplitude and death threshold. Constraints on these four parameters can be inferred from standard survival curves(*14*).

Here we show that variation in human lifespan is consistent with person-to-person differences in SR model parameters, subject to a strong constraint. Differences in damage production or removal rates greater than a few percent produce unrealistically long lifespans, whereas variation in threshold or noise preserves the observed upper limit near 120 years. This pattern is supported by analyses of NHANES(*16*, *17*) exposure cohorts, centenarian sibling data, ages of the longest-lived individuals, and historical cohorts adjusted for extrinsic mortality. As a contrasting case, survival curves from Hutchinson-Gilford progeria—a disorder of accelerated aging due to nuclear lamina defects—indicate altered production dynamics. Finally, we extended our analysis to additional mathematical models of aging and mortality and show that similar constraints apply. Together, these findings suggest that damage production and removal parameters in humans are tightly constrained with little person-to-person differences. Extending maximal human lifespan will require modifying the production or removal of aging-related damage, processes that appear largely unaffected by lifestyle, historical improvements, or common genetic variation.

## Results

### The saturating-removal model connects changes in the shape of the survival curve to damage dynamic parameters

The saturating-removal (SR) model mathematically describes aging and mortality dynamics across species (*10*, *13*, *14*, *18*), formulated as a stochastic differential equation (Fig. 1a). Central to the model is the simplifying assumption of a dominant, causal form of damage (X), that is upstream of the diverse and complex age-related changes such as inflammation and stem cell exhaustion. The SR model remains agnostic to the molecular identity of X, allowing applicability across organisms with distinct aging mechanisms.

**Fig 1.**
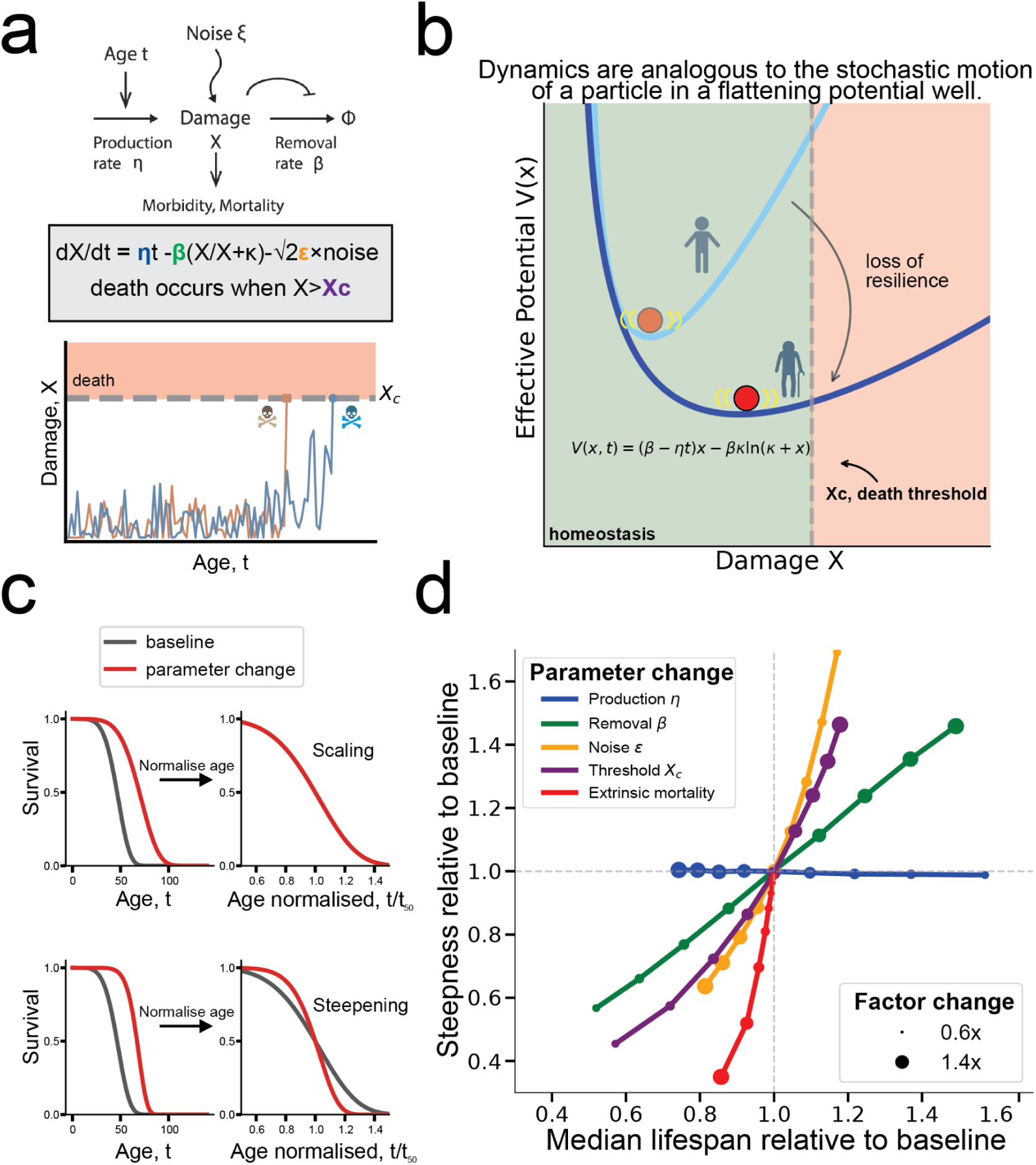
Threshold and noise parameters in the SR model primarily steepen the survival curve with mild effect on median lifespan. **a.** The saturating-removal model describes the stochastic dynamics of damage X that accumulates with age due to rising production and saturating removal. Simulations (orange and blue) show two individual stochastic trajectories crossing the death threshold Xc at different times **b.** SR model dynamics can be visualized as a particle jiggled by noise in a potential well that gradually flattens, reflecting decreasing resilience. The light and dark blue curves show the potential at young and old age. The threshold Xc marks the death threshold. **c.** Illustration of scaling and steepening effects upon change of parameter. **d.** Effect of SR parameter changes on median lifespan and steepness, defined as median lifespan divided by the interquartile range of the survival curve (Methods), relative to baseline. Marker size indicates fold-change of the parameter relative to baseline in increments of 0.1. Extrinsic mortality is zero at baseline and changes along the red curve from 10⁻⁴ year^-1^ to 10^-2^ year^-1^ in 10 logspaced steps. Control parameters η = 0.5 year^-2^, β = 54.7 year^-1^, κ = 0.5, ε = 51.8 year^-1^, Xc = 17.

In the SR model, damage X accumulates with age, produced by damage-producing units that accumulate linearly with age (production = ηt). Removal rate of damage saturates at high damage levels, modeled by Michaelis-Menten kinetics where removal = βX/(κ+X). Gaussian white noise ξ with amplitude ε represents stochastic fluctuations. Death occurs when damage crosses a threshold Xc (Fig. 1a).

Due to noise, the model generates stochastic, individual-specific damage trajectories and lifespans. The model reproduces the Gompertz law seen in human mortality data, including late-life mortality slowdown(*9*, *10*). In a recent study, variations in mouse survival curves across interventions were tied to specific changes in SR model parameters(*13*), as different model parameters influence median lifespan and survival curve steepness in distinct ways (Fig. 1c).

One can thus infer model parameters from the shape of survival curves. To do so, we quantify steepness as the ratio of median lifespan to inter-quartile range of the survival curve, consistent with previous definitions(*19*) (see Methods for details on this definition). Lowering the damage production rate parameter η extends both median lifespan without affecting steepness - a phenomenon known as scaling(*20*). Raising the removal rate β increases median lifespan and steepness proportionally. Changes in the death threshold (Xc) or noise amplitude (ε) modestly impact median lifespan and strongly impact survival curve steepness. These effects are summarized in a steepness-longevity plot (Fig. 1d), showing changes in median lifespan and steepness relative to baseline, when each parameter is changed. Baseline control parameters were fit to Swedish period mortality data from 2009(*10*).

We also incorporate extrinsic mortality into the SR model as a constant probability of death per unit time from external causes throughout life(*19*, *21*) (Methods). Extrinsic mortality sharply reduces steepness but has a small effect on median lifespan, except at high values (>5×10⁻³ year⁻¹). We also tested extrinsic mortality that rises with age based on human demographic data(*15*), and found nearly identical results to those obtained with constant extrinsic mortality (not shown).

In physics terms, the SR model is a Langevin equation for an overdamped particle trapped in a potential well. Death is modeled as a first-passage event, when the particle stochastically “crawls up” the potential well to cross a death threshold. Over time, the potential well flattens, reflecting a loss of resilience(*22*), increased sensitivity to fluctuations, and thus an increased chance of death (Fig. 1b). The shape of the potential well is determined by production η and removal β parameters, while the noise amplitude ε (analogous to temperature in statistical mechanics) quantifies the strength of fluctuations.

This framework prompts the question: which parameters vary across a human population? Do individuals differ primarily in the dynamics of their potential wells, the amplitude of noise, or their death thresholds? This is the question we seek to answer.

### Variations in threshold and noise, but not production or removal, preserve late-life survival statistics

Genetic and lifestyle differences affect demographic survival patterns, and therefore they should translate to variations in SR model parameters between individuals. This parameter heterogeneity, however, is constrained by two empirical observations:

1. Late-life survival curves (reliable data up to age ∼105) decline super-exponentially, and thus lifespans above 110 are exceedingly rare(*23*) (<0.1% of those reaching 100) (Fig 2a).
2. Mortality curves (describing chance of death per year) of different cohorts converge at advanced ages^24–26^, a phenomenon called the “compensation law of mortality”.

**Fig 2.**
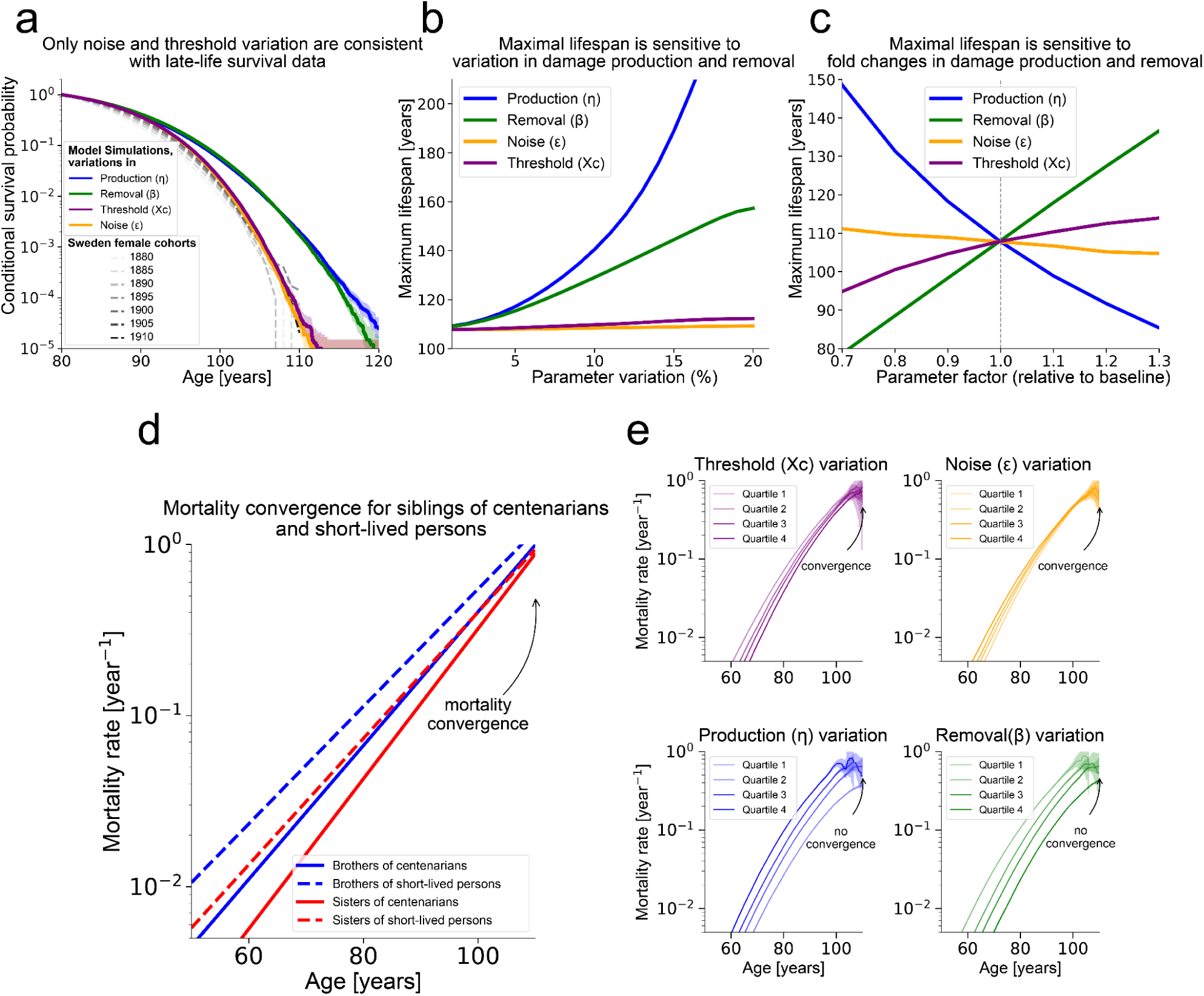
Variations in noise or threshold, but not production or removal, are consistent with late-life survival curves, maximal lifespan constraints, and mortality convergence. **a.** Late-life survival declines super-exponentially with age. Introducing 5% parameter heterogeneity shows that only variations in threshold (Xc) and noise (ε) maintain this steep decline, whereas equivalent heterogeneity in production (η) or removal (β) flattens it. Grey lines - mortality data for Swedish female cohorts. **b.** A proxy for maximum lifespan (age when survival = 10⁻⁴) under varying levels of normally-distributed parameter heterogeneity. Modest heterogeneity in production (η) or removal (β) extends lifespan beyond realistic limits, whereas variations in threshold (Xc) or noise (ε) preserve maximal lifespan. **c.** Effect of baseline parameter values on maximal lifespan. Changing baseline threshold or noise modestly affects lifespan, whereas changes in η or β substantially alter it. **d.** Best-fit Gompertz mortality curves from Gavrilova et al.(*24*), comparing siblings of centenarians (solid lines: blue, brothers; red, sisters) with siblings of short-lived individuals (dashed lines: blue, brothers; red, sisters) for U.S. cohorts born 1890-1897. **e.** Mortality curves for simulations with 5% heterogeneity in each parameter, stratified by quartiles. Only heterogeneity in threshold (Xc, upper-left) and noise (ε, upper-right) reproduce observed late-life mortality convergence. Baseline parameters in all panels: η = 0.43 year^-2^, β = 65.7 year^-1^, κ = 0.5, ε = 51.83 year⁻¹, Xc = 18, fit to Swedish female cohorts (1890-1900).

We asked which parameters can vary and obey these two constraints. To do so, we incorporate inter-individual variation by sampling model parameters for each simulated individual from a normal distribution, consistent with additive polygenic traits(*27*). Nearly identical results are found with lognormal distributions (not shown).

We find that even modest heterogeneity (>5% CV) in damage production rate η or removal rate β flattens late-life mortality, leading to unrealistically long lifespans (Fig. 2a,b, Supplementary Figure 2). In contrast, threshold Xc and noise amplitude ε can vary by more than 20% between individuals without disrupting late-life survival patterns (Fig. 2a, b). We recently showed(*15*) that ∼20% variation in threshold Xc reproduces lifespan correlations and survival statistics observed in monozygotic and dizygotic twin studies. Similarly, changes in the baseline parameters for the threshold (Xc) and noise (ε) have a very mild effect on maximum lifespan, whereas changes in production (η) and removal (β) alter it substantially (Fig. 2c). In the Supplementary Information, we provide the mathematical results behind these findings.

We also examined late-life mortality convergence. Mortality rates of long-lived and short-lived subpopulations are known to converge at very old ages - for example, among siblings of centenarians and siblings of individuals who lived only to age 65(*24*) (Fig. 2d). This convergence indicates that even siblings of centenarians, who likely possess favorable genetic factors for longevity, exhibit mortality patterns at advanced ages similar to those with less favorable genetics.

Introducing variation in threshold Xc, noise ε, production η, and removal β reveals differences in mortality convergence. Only variations in threshold or noise produce old-age convergence similar to empirical observations (Fig. 2e). In contrast, variations in damage production η or removal β create sub-cohorts with diverging or parallel survival curves that fail to converge at old age.

The intuitive reason for the unrealistic lifespans when production eta and removal beta vary is that these parameters create subpopulations that are very long-lived and dominate the extremes of the lifespan distribution. Both threshold and noise affect intercept and slope in compensating ways to cause convergence, capturing the inverse correlation between slope and intercept in human cohorts known as the Strehler-Mildvan correlation(*28*). In the Supplementary Information, we provide mathematical results behind this.

We conclude that damage production and removal rates are likely conserved across humans, varying at most by only a few percent. Larger variation would produce far more long-lived individuals than observed. Therefore, the parameters most likely to vary between individuals without distorting survival curves are threshold Xc and noise amplitude ε.

### Human cohorts with specific lifestyle exposures show changes consistent with threshold or noise

To test whether threshold and noise are the primary parameters that vary across human populations, we analyzed data from the National Health and Nutrition Examination Survey(*16*) (NHANES). This is a cross-sectional survey conducted by the U.S. Centers for Disease Control and Prevention to assess the health and nutrition of the U.S. population.

We stratified the NHANES cohorts based on self-reported lifestyle exposures with well-established effects on longevity. These included diet quality^29–32^, income level(*33*, *34*), alcohol consumption^35–37^, physical activity^38–40^, sleep duration and quality^40–48^, social support^49–51^, church attendance(*52*, *53*), and education level(*54*, *55*) (Methods). We linked these cohorts to mortality data updated through December 31, 2019(*17*), pooling females and males together.

For each exposure subgroup, we calculated median lifespan and survival curve steepness, comparing them to values for the full cohort (n = 60,000). To adjust for extrinsic mortality, we fit a Makeham-Gamma-Gompertz (MGG) model to the total cohort mortality curve (Methods), and then set the Makeham term to zero (m_ex_ = 0). We tested whether each subgroup deviated significantly from this adjusted baseline in terms of median lifespan and steepness. This allowed us to assess whether observed differences were due to intrinsic parameter changes or elevated extrinsic mortality.

We plotted each group’s steepness and longevity, normalized to the total cohort values on a steepness-longevity plot (Fig. 3), similar to Fig. 1c. Exposure group cohorts were not adjusted for covariates, as our aim was not to estimate hazard ratios for specific exposures (as in a Cox regression), but rather to explore directional variations in the steepness-longevity space across exposure groups. Specifically, we expected that differences between exposure cohorts should correspond with changes in threshold (Xc) or noise (ε).

**Fig 3.**
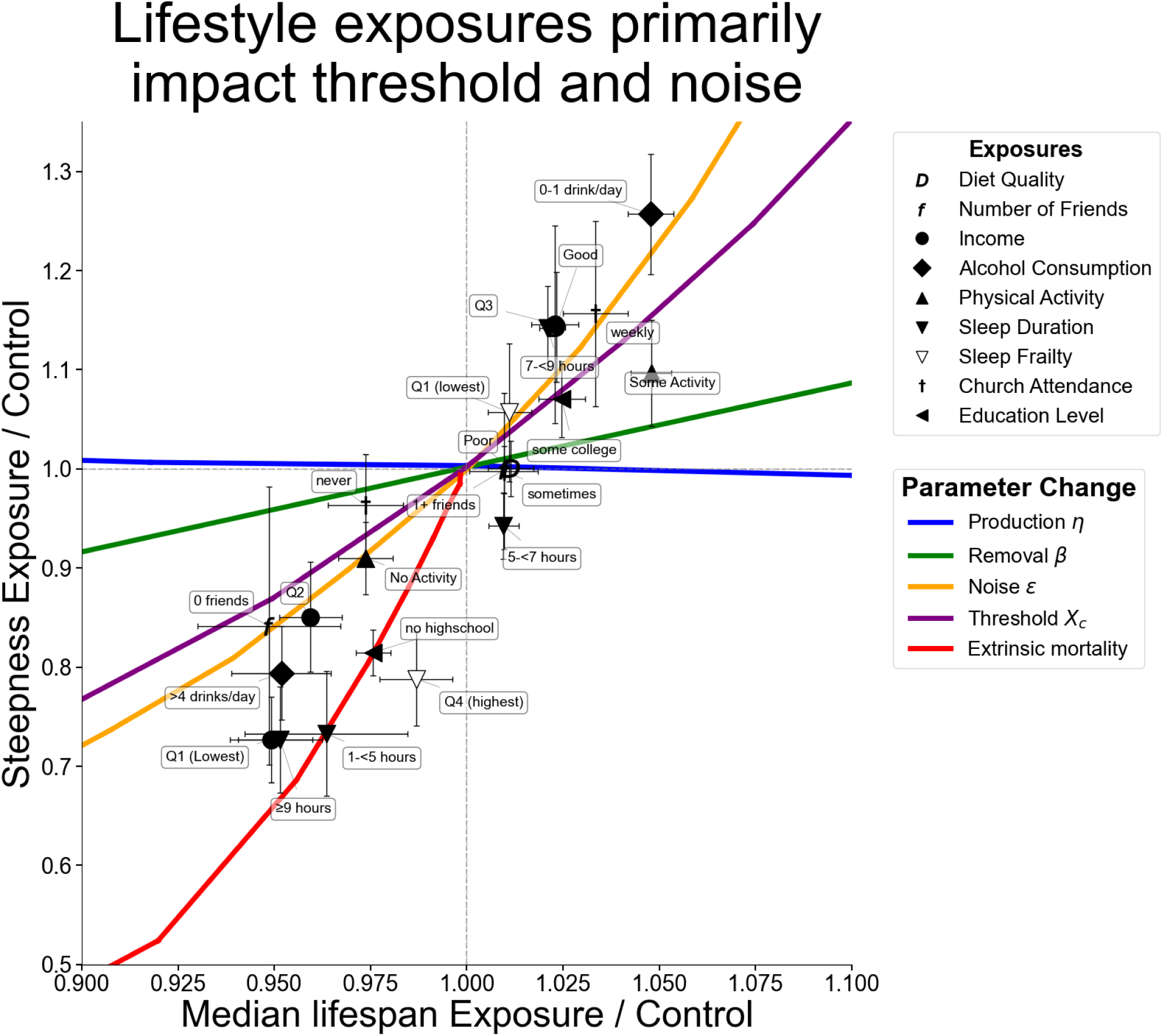
Median lifespan and steepness for lifestyle exposure cohorts from NHANES, relative to the total cohort. Cohorts cluster along trajectories consistent with changes in threshold (Xc) and noise (ε). Lines indicate predicted trajectories for various fold-changes in SR model parameters. All survival curves (both NHANES and SR model) are conditioned on survival to age 20. Baseline SR parameters were fitted to U.S. 2019 mortality data (η = 0.54 year^-2^, β = 54.75 year⁻¹, κ = 0.5, ε = 51.83 year⁻¹, Xc = 21) with 20% heterogeneity in Xc, based on previous analysis of twin studies(*15*). Exposure group definitions, sample sizes, and deaths per group are detailed in the Methods and Supplementary Information. Error bars represent bootstrap-derived standard errors.

We find that exposure groups associated with advantageous longevity exhibited a substantial rise in steepness (10-30%) alongside modest gains in median lifespan (1-5%). These cohorts included alcohol consumption of 0-1 drinks/day, sleep duration of 7-9 hours, good diet quality, high income, weekly church attendance, low sleep frailty (Methods), physical activity, and college education. These groups clustered along the noise and threshold curves in the steepness-longevity plot, consistent with a reduction in noise ε or gain in threshold Xc. However, changes in production η and removal β parameters are generally ruled out.

These observations are consistent with prior findings linking beneficial longevity outcomes to adequate (but not excessive) sleep duration(*41*, *45*, *48*), protective effect of moderate alcohol intake(*35*, *36*), high-quality diet(*30*, *31*), social support networks^49,51^, regular religious or community participation(*52*, *53*), physical activity(*38*), higher education level(*54*), and economic stability(*33*).

Among shorter-lived groups, some clustered along the extrinsic mortality curve, such as cohorts with no high school education, <5 or >9 hours of sleep, lowest sleep-quality quartile, lowest income quartile. These patterns align with evidence that sleep deprivation raises the risk of accidents and infections(*42*, *57*), and that low-income and low-education groups face elevated risks from violence and deaths of despair^33,58–61^.

Additional short-lived exposure cohorts appeared to primarily elevate noise or reduce threshold. These included the second income quartile, consuming >4 drinks/day, lacking close friends, no physical activity, or not attending church. These align with increased mortality risks associated with excessive alcohol intake^35–37^, social isolation(*51*), and sedentary lifestyle(*38*, *39*).

### Historical cohorts show changes in threshold and noise

We also analyzed the parameters underlying historical cohorts. Since the mid-19th century, both median lifespan and lifespan equality have increased(*4*, *6*) - a trend referred to as the rectangularization of the survival curve. These improvements are largely attributed to reductions in childhood mortality and other forms of extrinsic mortality, driven by advances in sanitation, disease control, healthcare access, and related factors(*62*, *63*). We sought to determine whether historical gains in longevity reflect only decreases in extrinsic mortality or also involve changes in other model parameters.

To test this, we analyzed Swedish and Danish period survival curves from 1800 to 2020 from the Human Mortality Database (HMD)(*64*), tracking changes in steepness and median lifespan (Fig. 4a). We plot the historical changes for Swedish males and females combined with all values normalized to the year 2020 curve.

**Fig 4.**
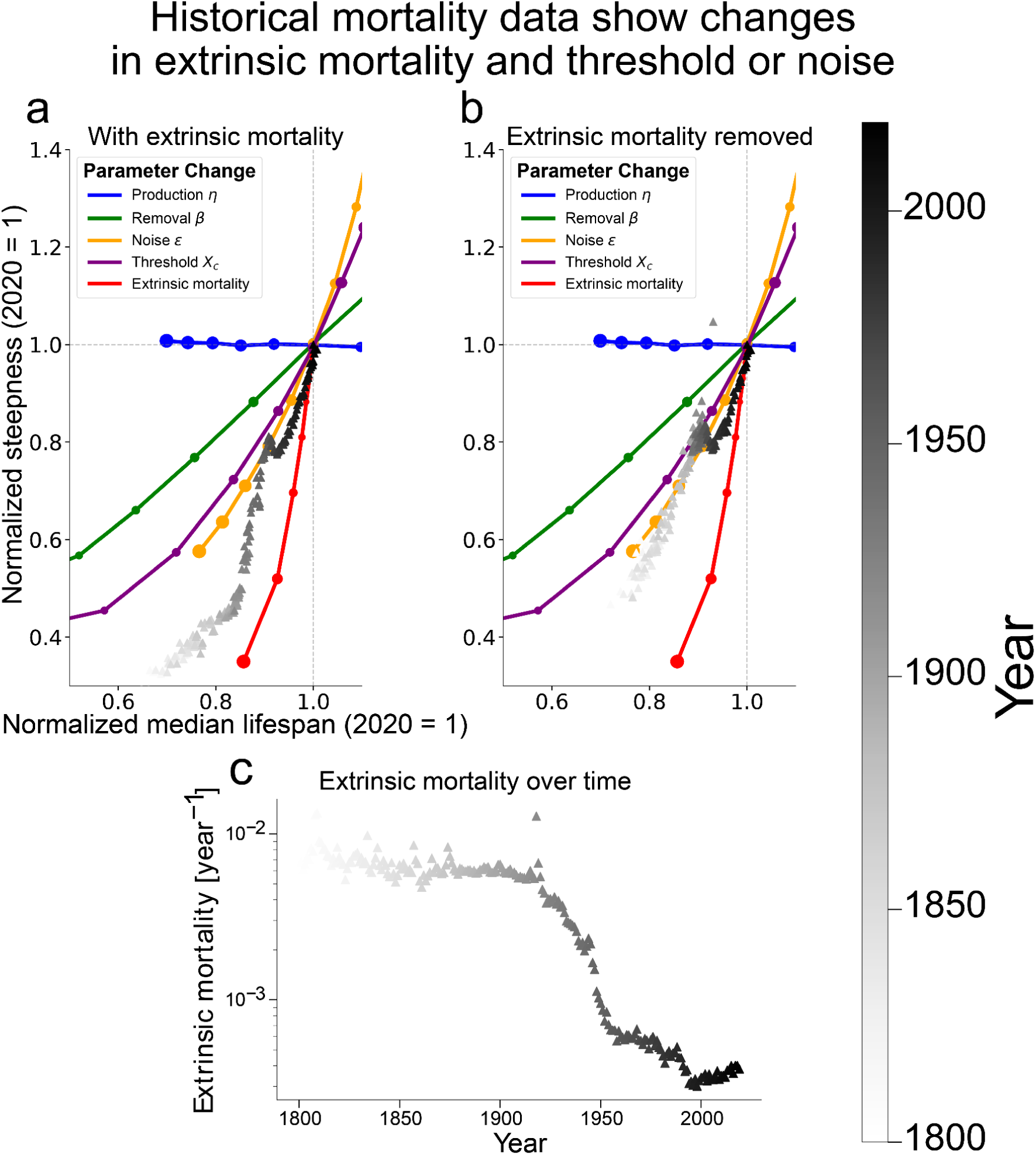
Historical mortality improvements after adjusting for extrinsic mortality are driven by changes in threshold and noise. **a.** Steepness-longevity plot for Swedish period survival curves (males and females pooled), 1800-2000 (shaded by year), normalized to year 2020 data (from HMD(*64*)). Lines show predicted trajectories for ±70% changes in model parameters. Baseline parameters fitted to Sweden 2019 data: η = 0.49 year^-2^, β = 54.8 year⁻¹, κ = 0.5, ε = 51.8 year⁻¹, Xc = 17. **b.** Same analysis after removing extrinsic mortality (setting m_ex_=0 in the Makeham-Gamma-Gompertz fit). **c.** Extrinsic mortality parameter m (log scale) fitted to Swedish period data (males and females pooled).

In the steepness-longevity plot (Fig. 4a), historical changes initially follow the noise/threshold curves, then rise sharply in steepness, driven by a nearly tenfold drop in extrinsic mortality between 1900 and 1950 (Fig. 4c). This decline reduced early deaths and steepened the survival curve.

Given the strong influence of extrinsic mortality on these trajectories, we sought to adjust for extrinsic mortality in order to reveal underlying parameter shifts. We removed the Makeham extrinsic mortality term from each cohort (Methods) and recalculated steepness and lifespan (Fig. 4b). After removing extrinsic mortality, the change of survival curve shapes is consistent with a ∼35% increase in threshold or a ∼60% increase in noise over this historical period (Fig. 4b). Changes in production or removal parameters do not fit the data. Danish survival curves show similar trends (Supplementary Fig. 1).

There is a phase (∼1940-1970) where median lifespan increases while steepness declines (Fig. 4a,b), a pattern previously noted(*65*, *66*). This behavior is not captured by any single SR model parameter change. This effect was suggested to arise when gains in survival are concentrated at older ages while underlying disparities in health or risk persist across the population(*65*, *66*).

In sum, after accounting for extrinsic mortality, historical changes in survival curve shape from 1800 to 2020 are largely explained by reduced noise and/or increased threshold.

### Modest increase in recorded maximal human lifespan is captured by changes in threshold

The age of the oldest human alive in a given year increased from 109 in 1960 to 117 in 2020(*67*) (Fig. 5a), following an approximately linear trend. Similar linear trends are seen for different ranks (2nd oldest, 10th oldest etc) (Supplementary Figure 5). We asked what drives this linear rise. Three possibilities were considered: (1) that the maximal age increased due to population growth, (2) that it grew due to decreased extrinsic mortality, or (3) that it reflected changes in SR model parameters.

**Fig 5.**
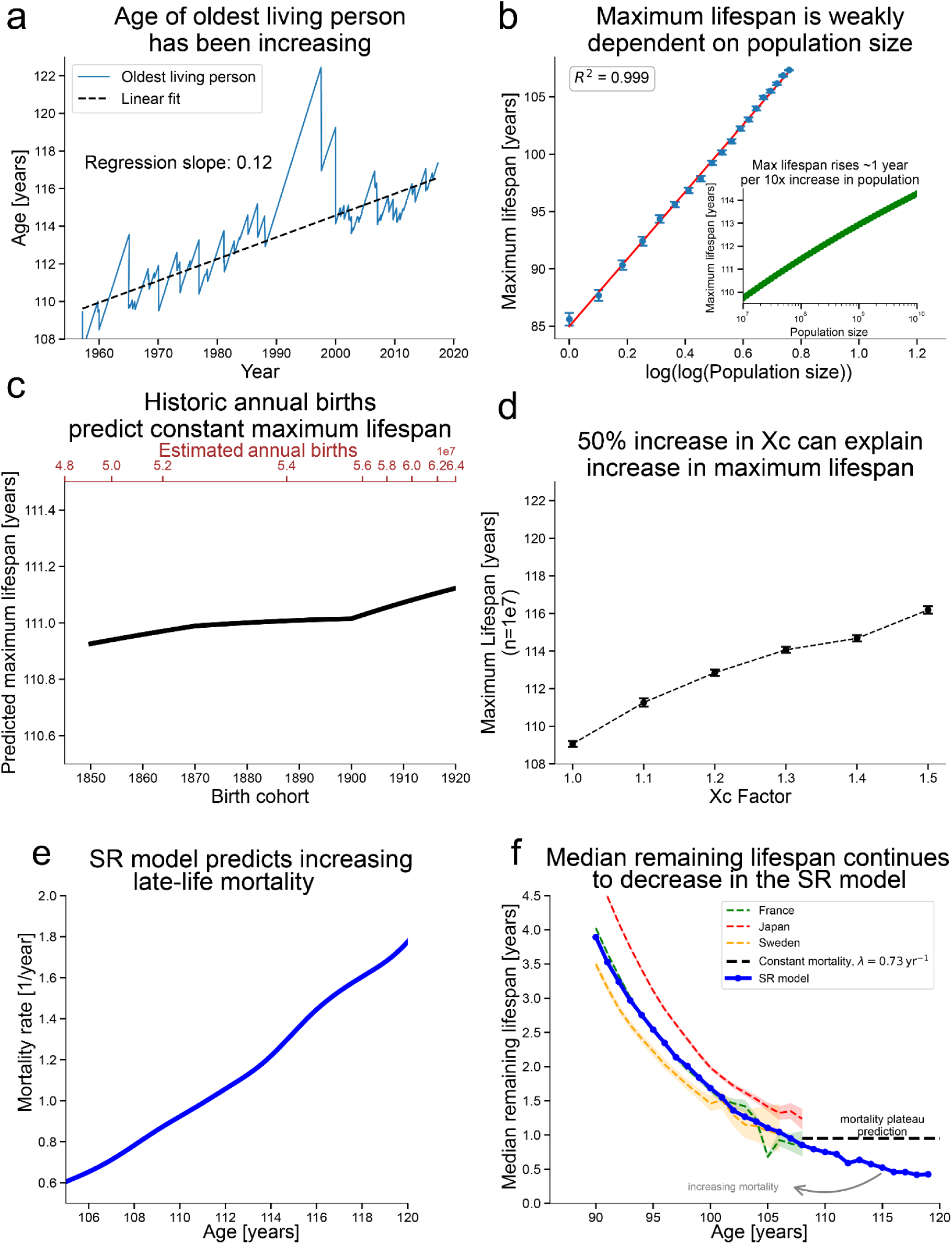
Increase in age of the oldest living person can be explained by a 50% rise in threshold (Xc), but not by population growth. **a.** Observed age of the oldest living person, data from the Gerontology Research Group(*67*, *72*). The dashed black line indicates the linear trend in record age, with a slope of 0.12. **b.** Maximum lifespan (blue dots) in SR simulations with different cohort sizes. Error bars are standard errors. The red line is the best-fit linear regression. Inset: Same as (b), but with population size on a logarithmic scale. **c.** Predicted maximum lifespan based on the regression in (b), using estimated annual global births from 1850-1920. This period roughly corresponds to the birth cohorts of the oldest living individuals shown in (a). **d.** Effect of increasing the threshold Xc on simulated maximum lifespan for n = 10⁷ individuals - approximately one quarter of annual global births during 1850-1920 - representing industrialized countries. **e.** The mortality rate among supercentenarians rises approximately linearly in the SR model. The simulation shown uses a 1.5× scaling of the baseline threshold parameter Xc. **f.** Median remaining lifespan estimates for France, Japan, and Sweden (period 2020) are shown, based on observed exposure and death rates, together with predictions from the SR model. For reference, the dotted black curve corresponds to a constant mortality rate of λ = 0. 73 *year*^−1^. The simulation again uses a 1.5× scaling of Xc relative to baseline. Baseline parameters for all panels: η = 0.66 year^-2^, β = 62.05 year⁻¹, κ = 0.5, ε = 51.83 year⁻¹, Xc = 14, with 20% person-person variation in threshold Xc.

To test the population-size effect, we simulated the SR model and found that maximal lifespan scales weakly as log(log(N)), where N is the number of simulated individuals (Fig. 5b). From mathematical considerations, the slope between maximal lifespan and log(log(N)) should be roughly equal to the inverse of the Gompertz exponent, assuming Gompertzian mortality (see Supplementary Information). This roughly corresponds to an increase of roughly one year in maximal lifespan for every tenfold increase in population size in the historical range (Fig. 5b, inset). The historic growth in global annual births(*68*) could not account for the observed rise, and instead predict a nearly constant predicted maximal lifespan given constant model parameters (Fig 5c). We conclude that population growth during this period does not explain the increase in maximal age.

Extrinsic mortality - deaths caused by external factors such as infections, accidents or violence(*19*, *21*) - has a negligible impact on maximum lifespan. Even under historically high extrinsic mortality rates observed in the early 19th century (around 10⁻² year⁻¹), the fraction of total deaths attributable to these causes would be less than 0.5. Under the most extreme assumptions, eliminating extrinsic mortality would therefore be roughly equivalent to doubling the effective population size. However, because maximum lifespan scales only weakly with population size through a log(log N) relationship, such a change would not meaningfully alter the observed upper limit of lifespan.

After ruling out population growth and extrinsic mortality, we asked whether temporal changes in model parameters, specifically in the threshold Xc, could account for slow rise in maximal lifespan. To characterise the supercentenarian tail of survival in the SR model, we use a constant-population “go with the winners” algorithm that repeatedly clones surviving trajectories (Methods). Keeping cohort size fixed at n=10⁷ (same order of magnitude as annual births in the relevant period, Methods) - we varied Xc over time. We found that a 50% rise in Xc over the relevant period reproduces the observed linear rise in maximal lifespan (Fig 5d), including the top fifty recorded deaths in that calendar year (Supplementary Fig. 5). We thus conclude that historical increases in Xc can explain the modest increase in human maximal lifespan.

The SR model offers a prediction for whether supercentenarian mortality continues rising at very late ages, or instead levels off. Recent studies argue that mortality for supercentenarians is consistent with a plateau of roughly 0.73 per year(*69–71*). In the SR model, mortality keeps increasing, though its rate of growth slows substantially at extreme ages (Fig. 5f). Between 110 and 120 years, the model produces an approximately linear rise. In the asymptotic limit, mortality is expected to grow quadratically with time rather than converge to a constant value (see Supplementary Information). This continued increase implies a declining mean remaining lifespan with age, in contrast to the constant expectation time that would follow from a memoryless, constant-mortality process (Fig. 5f). Future data at extreme ages can test this prediction more rigorously.

### A disease of accelerated aging cannot be explained by changes in threshold and noise alone

After showing that genetic variation, lifestyle, and historical factors influencing longevity are consistent with changes in threshold or noise, we asked whether accelerated aging diseases are also due to changes in these parameters. To test this, we analyzed the largest published survival dataset of untreated patients with Hutchinson-Gilford Progeria Syndrome (HGPS) (n = 202)(*73*).

HGPS is a rare genetic disorder occurring in roughly 1 in 1-4 million live births, caused by a mutation in the *LMNA* gene(*74*). This mutation results in the production of progerin, a truncated form of lamin A, which compromises the integrity of the nuclear membrane. This disruption leads to accelerated aging symptoms in children with changes similar to natural aging(*75*, *76*) but occurring about 6-fold faster, including growth failure, early-onset cardiovascular disease, and premature death (median lifespan ∼14 years)(*74*).

In Fig. 6a, we plot the survival curve of HGPS patients alongside period survival curves from 2019 USA and Sweden, normalized by median lifespan. Compared to normal populations, the HGPS curve is >50% shallower, with a tail reaching nearly twice the median lifespan. This shape is not seen in any lifestyle exposure group or historical cohort (such shallowness would mean that individuals would live to 160 if the median is 80).

**Fig 6.**
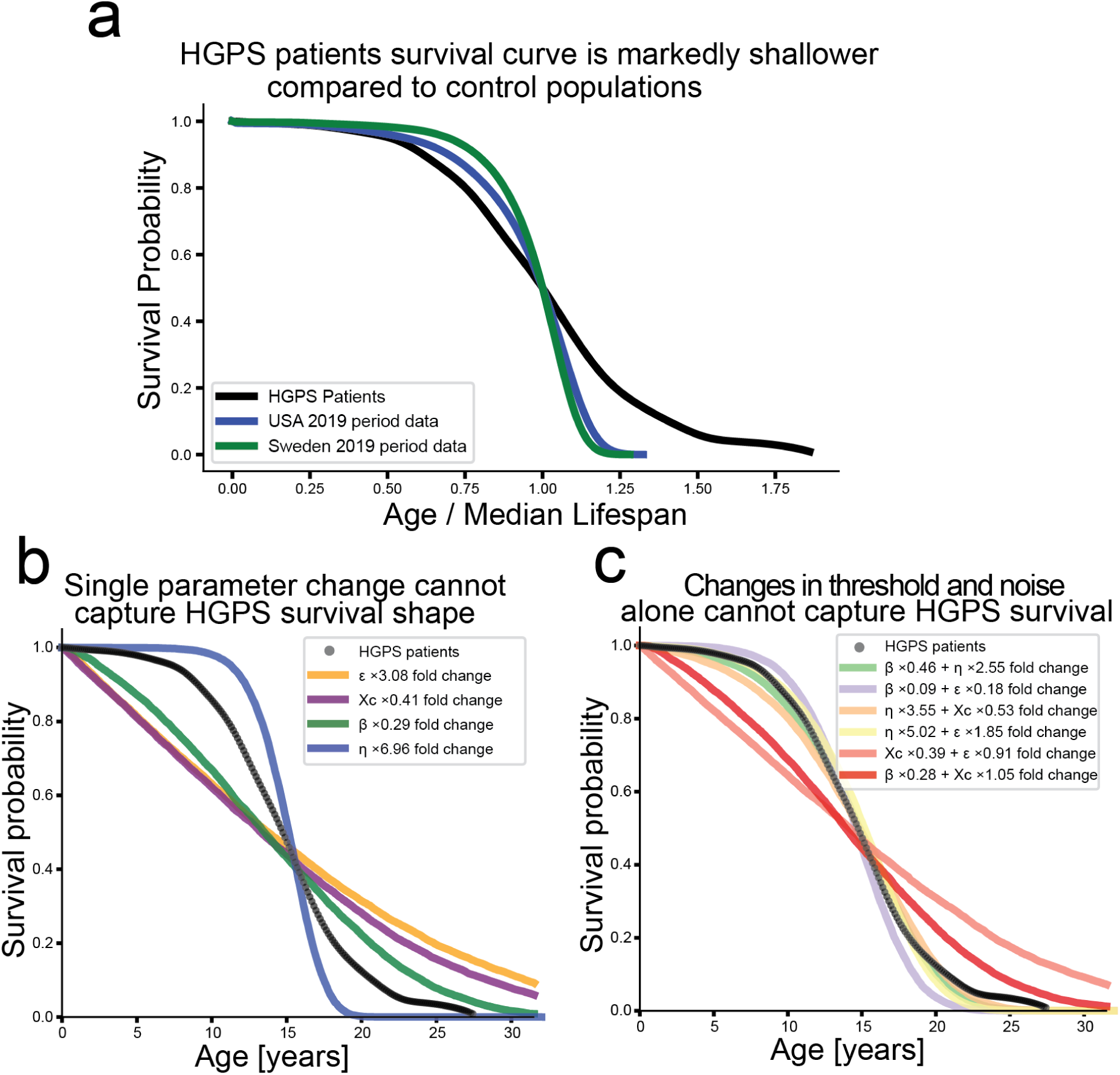
Progeria survival curves can not be explained by changes in threshold and noise, and require increased production or reduced removal. **a.** Survival curve for HGPS patients (black, n = 202; data from Gordon et al.(*73*)) normalized by median lifespan, shown alongside 2019 period survival curves for the U.S. (blue) and Sweden (green), (males and females pooled). **b**. Best-fit SR model allowing a single parameter to vary relative to baseline. No single parameter change captures HGPS survival curve shape. **c**. Best-fit SR model allowing two parameters to vary simultaneously. Fitting the HGPS curve requires changes in production (η) or removal (β); changes in both threshold (Xc) and noise (ε) cannot reproduce the curve on their own. Fold changes are relative to baseline parameters fitted to Sweden 2019 data: η = 0.49 year⁻^2^, β = 54.8 year⁻¹, κ = 0.5, ε = 51.8 year⁻¹, Xc = 17.

We asked which changes in the SR model parameters could reproduce the HGPS survival curve. Because the changes in both steepness and median lifespan exceed twofold for HGPS, the position within the steepness-longevity plane falls outside the linear regime and cannot be interpreted directly. We therefore fitted the full survival curve under different parameter changes. Varying each parameter individually showed that no single parameter could fit the data (Fig. 6b).

Allowing two parameters to change simultaneously revealed that changes in threshold (Xc) and noise (ε) together cannot reproduce the survival curve on their own (Fig. 6c). In contrast, increasing damage production (η) by a factor of 2-4 or decreasing removal (β) by a factor of 2-3, together with variation in a second parameter, recapitulates the observed survival pattern.

Based on biological evidence, HGPS primarily affects nuclear structure and DNA maintenance, leading to cellular senescence(*74*), consistent with an increase in production rate η. However, a reduction in removal β cannot be ruled out, potentially reflecting vascular damage and immune dysfunction. We conclude that this severe premature-aging disorder cannot be explained by changes in threshold or noise (Xc or ε) alone, but instead requires substantial alterations in production or removal - parameters that we find to be largely insensitive to lifestyle or environmental exposures.

### Other models of aging exhibit similar parameter constraints

We tested two additional mathematical models of aging to evaluate whether our results are model-dependent. We focused on how parameter variation affects maximal lifespan and how baseline parameter shifts map onto the steepness-longevity plane.

First, we tested the Makeham-Gompertz equation, a commonly used three-parameter fit for human mortality curves that has no mechanistic basis, defined as *m*(*t*) = *ae*^*bt*^ + *m*^*ex*^. We found that varying slope or intercept alone produces implausible maximal lifespans. In contrast, coupled changes in slope and log-intercept that preserve the Strehler-Mildvan correlation(*28*) (*log*(*a*) ∝ *b*) leave maximal lifespan unchanged (Fig. 7a, see the Supplementary Information for the mathematical reasoning for this). These correlated shifts also generate the correct trajectory in the steepness-longevity plane seen in historical data and exposure groups (Fig. 7b). Thus, realistic heterogeneity in Gompertz curves requires maintaining the Strehler-Mildvan relation. The SR model naturally produces this correlation through variation in Xc or ε (see Supplementary Information).

**Fig 7.**
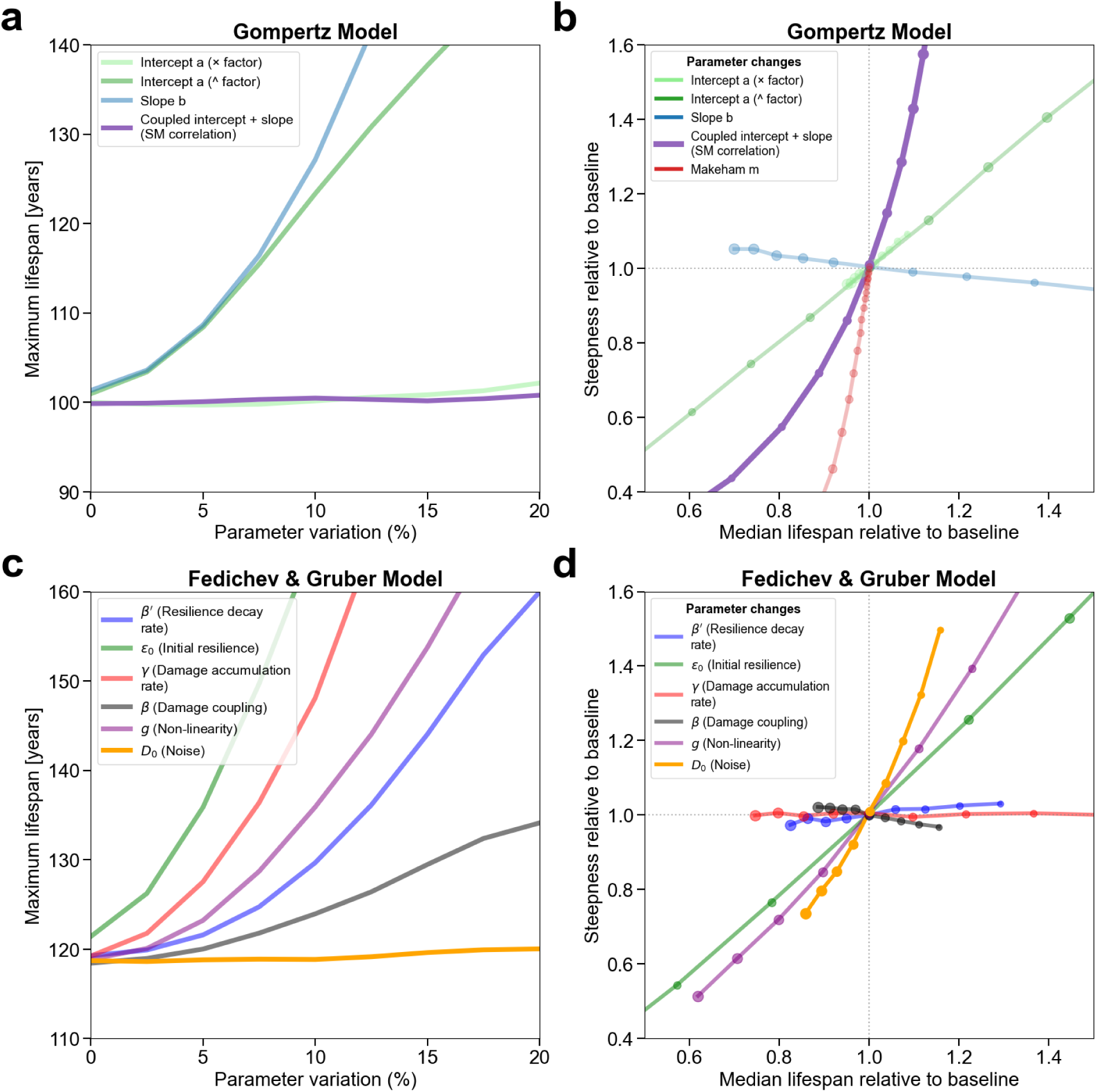
Analysis of maximum lifespan and steepness-longevity in the Gompertz and Fedichev & Gruber (FG) models. **a.** Maximal lifespan (age at survival = 10^-4) under varying degrees of parameter heterogeneity in the Gompertz-Makeham model. Only correlated variation in slope and log(intercept) preserves maximal lifespan. **b.** Steepness-longevity plot (as in Fig. 1c), showing effects of baseline parameter shifts in the Gompertz model. Marker size indicates fold-change in parameter values from 0.6 to 1.4 in increments of 0.1. **c.** Same as a but for the FG model. **d.** Same as b but for the FG model. Baseline parameters for the Gompertz-Makeham model: a = 2e-5 year^-1, b = 0.1 year^-1, m_ex = 0. Baseline parameters for the FG model: β′ = 0.013, ε₀ = 4, γ = 1, β = 0.015, g = 0.8, D₀ = 1.1.

We next analyzed the recent Fedichev & Gruber (FG) minimal model of aging(*77*). Similar to the SR model, the FG model is an equation for the stochastic dynamics of a primary health variable, and can also be visualized as a particle jiggling within a flattening potential well (see Methods, Supplementary Information for a mathematical analysis of the model’s predictions). Unlike the SR model however, the FG model has no threshold parameter governing death. Instead, at a critical time the model undergoes a saddle-node bifurcation when the protective potential barrier collapses, imposing a strict maximal lifespan limit. The SR model differs in allowing survival beyond the stability point due to its threshold parameter.

We tested how variation in each parameter impacts maximal lifespan and survival-curve shapes. Variation in noise amplitude respects maximal lifespan constraints, similar to the SR model, whereas changes in dynamic rate parameters yield unrealistic lifespans (Fig. 7c). The nonlinear coupling parameter g - analogous to the SR model’s Xc - also influences maximal lifespan, highlighting a structural difference between the models. Only noise variation produces realistic steepness-longevity curves matching empirical data (Fig. 7d).

Our results point to a core constraint in mechanistic aging models: the parameters that control the time-dependent flattening of the potential well must be tightly conserved in humans. In the SR model these are η and β; in the FG model they are γ, β, β′, and ε_0_. These dynamic parameters set the pace at which stability is lost, and if they were substantially varied across individuals we would see late-life mortality flattening and unrealistically long lifespans. Observable heterogeneity instead aligns with static parameters like threshold or noise amplitudes. Such variation naturally preserves the Strehler-Mildvan correlation (see Supplementary Information).

## Discussion

We explored the question of maximal human lifespan using the saturating removal model of damage dynamics. Only person-to-person variation of at most a few percent in damage production (η) or removal (β) is compatible with late-life survival; larger variation predicts too many extremely long-lived individuals. In contrast, broad variation in threshold (Xc) or noise (ε) respects late-life survival patterns and preserves mortality convergence. Analyses of NHANES exposure cohorts and historical survival data indicate that lifespan gains are driven primarily by higher threshold and lower noise - reflecting greater physiological robustness and reduced stochastic variability - whereas damage production η and removal β remain essentially unchanged. Common genetic and lifestyle factors seem to modulate threshold and noise rather than damage production or removal. This constancy of damage production rate is striking given that damage production varies by roughly seven orders-of-magnitude across species, from C. elegans to humans(*14*), suggesting that long-term evolutionary shifts in lifespan act largely through changes in damage production.

This distinction between conserved and variable parameters extends beyond the SR model. Our analysis of the Fedichev & Gruber (FG) model confirms that parameters driving the time-dependent flattening of the stability landscape cannot vary substantially between individuals without violating demographic limits. Demographic variation is instead found in either threshold or the intensity of random perturbations.

Improvements in threshold can also explain the slow increase in the age of the oldest living human, which increased by roughly 7 years over the past half-century. Population growth alone cannot account for this trend, as it predicts at most a one-year rise in maximal lifespan for every order-of-magnitude increase in population size. The SR model predicts a slow rise in mortality at supercentenarian ages, producing a declining median remaining lifespan. Current estimates remain consistent with a plateau, but improved data at extreme ages will allow a clearer test of this prediction.

What mechanisms drive increases in threshold, the maximal damage compatible with life? Improved medical care, improved cardiovascular capacity, infection resistance, and reduced frailty all effectively raise Xc. Also, lowering constant, age-independent damage from toxins or pathogens also equivalently increases Xc, since adding a constant damage term is equivalent to lowering Xc.

In contrast, Hutchinson-Gilford progeria survival curves require altered damage production (η) or removal (β), consistent with elevated cellular stress, chromatin disruption, enhanced DNA damage caused by the LMNA mutation (higher η), or impaired clearance and immune deficiency from vascular dysfunction (lower β). Thus, accelerated aging necessarily involves changes in damage production or removal.

The SR model parameters have plausible biological counterparts. We interpret X as the burden of damaged or inflammatory cells (including senescent cells). The production rate η reflects the accumulation of long-lived “error factories” that produce damaged cells, such as epigenetically altered stem cells; the removal rate β captures immune and vascular clearance; Xc indicates organismal robustness; and ε represents short-term physiological fluctuations (circadian rhythm, stress, immune variability).(*14*) This explains why lifestyle predominantly affects Xc and ε, extending median but not maximal lifespan.

Our results suggest that extending maximal human lifespan will require interventions that reduce damage production (via cellular reprogramming, enhanced repair, mTOR/mitochondrial interventions) or increasing damage removal (via improved immune/vascular clearance, senolytics). In mice, such interventions extend both median and maximal lifespan(*13*), but in humans, clear evidence is lacking. Common genetic variants and lifestyle factors minimally influence η or β, but rare variants might, analogous to progeroid mutations. Genomic screens targeting these pathways - such as DNA repair variants linked to delayed menopause(*78*) - may thus identify longevity-associated alleles, providing avenues to significantly alter the human lifespan limit.

## Methods

### Saturating-removal model

We used the saturating-removal (SR) mathematical model(*10*) of aging dynamics, and expanded it here to simulate heterogeneous cohorts. The SR model is a stochastic differential equation that describes the dynamics of damage that is assumed to be causally linked to mortality. Its core assumption is that a dominant form of damage, denoted X(t), drives age-related organismal changes.

The SR model describes damage X(t) as a balance of production, removal, and noise

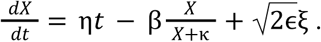

η, β, κ, ɛ are model parameters, while ξ represents Gaussian white noise. η has units of time⁻², β of time⁻¹, ε of time⁻¹, κ is normalized to 1, and X and Xc are in units of κ. The model is simulated using an Euler-Maruyama numerical scheme with n=10^6^ agents, with a reflecting boundary condition at *X* = 0. Death occurs upon crossing a critical threshold *X_C_*. Extrinsic mortality *m_ex_* is incorporated by adding a death probability of 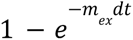 at each timestep, dt. Population heterogeneity was modeled by sampling model parameters from Gaussian distributions with specified means and standard deviations. Survival and mortality curves are computed from the lifespans generated for the simulated cohort.

Because the SR model allows only semi-analytical solutions, parameters were estimated by fitting survival curves using least-squares optimization (L-BFGS-B).

### Go-with-the-winners simulation

To probe the late-time survival tail in the SR model, we use a constant-population splitting algorithm of the “go-with-the-winners” type. The method evolves an ensemble of stochastic trajectories under the SR dynamics, where each particle carries its own parameter set. To preserve heterogeneity during resampling, particles are first grouped into bins in parameter space using quantile-based binning of the varying parameters. At each time step, all particles are advanced according to the SR model, and any particle that crosses its absorbing threshold Xc is replaced by a clone of a surviving particle from the same parameter bin. The cloning step uses weight splitting: if a survivor with weight w is chosen as a donor, then the donor and its clone are both assigned weight w/2. This construction maintains an unbiased representation of the killed SR model while reallocating computational effort toward trajectories that survive for long times, thereby enhancing resolution in the tail of the survival distribution. To control numerical underflow caused by repeated splitting, weights are periodically rescaled within each bin by a common factor, and the number of rescalings per bin is recorded and later used to undo this scaling when reconstructing the total survival probability; the mortality rate is computed at each time step from the ratio of the weighted mass that crosses the threshold to the weighted mass that remains below it. Lifespans are subsequently sampled from the survival curve, with bootstrapping for errors.

### Makeham-Gamma-Gompertz mortality model

The Makeham-Gamma-Gompertz model(*79*, *80*) is a parametric (non-mechanistic) model that describes the mortality of human data, given by

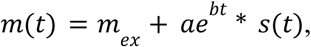

where *s*(*t*) accounts for late-life mortality deceleration, 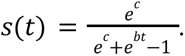 The Makeham-Gompertz model is recovered in the limit *c* → ∞. To model population heterogeneity, we implemented Gaussian parameter distributions in a scale parameter q such that *b* = *q b*_0_, *log*(*a*) = *log*(*a*_0_)/*q*. This approach ensures the Strehler-Mildvan correlation (*28*) between log intercept and slope is maintained and yields realistic maximal lifespan estimates. Population heterogeneity was modeled by sampling model parameters from Gaussian distributions with specified means and standard deviations. Lifespans for the MGG model were generated using inverse transform sampling. Survival and mortality curves are computed from the lifespans generated for the simulated cohort.

Model parameters were fitted to country death rates, log-averaged across all birth years in a cohort, using scipy’s curve_fit for non-linear least squares optimization. Fitting was performed in log-space to account for the exponential increase of hazard with age.

### Fedichev & Gruber’s minimal model of aging

Fedichev & Gruber’s minimal model of aging(*77*) describes the dynamics of the critical, slow mode of a composite health matrix, described by the following stochastic differential equations:

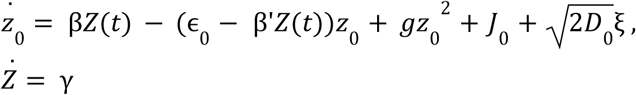

where ξ is Gaussian white noise. The model is simulated using an Euler-Maruyama numerical scheme, with n=10^5^ agents. Population heterogeneity was modeled by sampling model parameters from Gaussian distributions with specified means and standard deviations. Death is defined as the moment when the variable crosses the top of the protective potential barrier. The barrier location *Z^+^*_0_(*t*), is obtained by finding the roots of 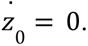 When introducing parameter heterogeneity, each agent has its own *Z^+^*_0_(*t*), based on its respective model parameters. Survival and mortality curves are computed from the lifespans generated for the simulated cohort.

### Steepness of survival curves

We define steepness as the median lifespan divided by the interquartile range of the survival curve: 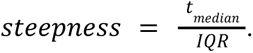 Other definitions are possible, such as using the inverse coefficient of variation µ/σ. However, this latter measure is heavily influenced by the distribution tail. In the exposure groups, too few deaths occurred to estimate the tail reliably, which makes tail-sensitive metrics unstable. Using 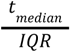 avoids this problem and provides a robust description of the survival curve’s shape.

### Extrinsic mortality

Extrinsic mortality for exposure groups and historical cohorts was estimated by fitting a Makeham-Gamma-Gompertz model (see above) to the respective mortality curve, and taking the fitted mex as the extrinsic component. Extrinsic mortality was removed by setting m_ex_ to 0.

### NHANES exposure groups

We analyzed publicly available questionnaire data from the National Health and Nutrition Examination Survey (NHANES), including only participants eligible for mortality linkage. Our analysis included 59,064 participants, of whom 9,249 (15.66%) died, with a median age at death of 82.1 years; the minimum entry age for participants was 18.0 years. We examined nine lifestyle and socioeconomic exposures in this study.

1. Diet Quality

- **Variable:** DBQ700 (“How healthy is your overall diet?”)
- **Cycles:** 2005-2018
- **Definition:** “Excellent”/“very good” responses (1-2) were categorized as “Good,” while “good”/“fair” responses (3-4) were categorized as “Poor.” Responses 5-6 and invalid codes (7, 9) were excluded.
- **Groups:** Good vs. Poor
2. Income-to-Poverty Ratio

- **Variable:** INDFMMPI (family monthly poverty level index)
- **Cycles:** 2007-2018
- **Definition:** Participants were divided into quartiles based on their family monthly poverty level.
- **Groups:** Q1 (lowest) to Q4 (highest)
3. Physical Activity

- **Variables:** PAQ605-PAD675 (vigorous/moderate work, transport, recreation)
- **Cycles:** 2007-2018
- **Computation:** MET-minutes/week were calculated across five domains: vigorous work (8.0 METs), moderate work (4.0), transport (4.0), vigorous recreation (8.0), and moderate recreation (4.0). Invalid responses (7, 9, 77, 99, 777, 999, 7777, 9999) were either excluded or replaced with 0 if they indicated “does not do this activity.” Totals were log-transformed [log(x+1)] and then min-max normalized (0-1).
- **Grouping:** Binary classification based on any reported activity.
- **Groups:** No activity (index = 0) vs. Some activity (index > 0)
4. Sleep Duration

- **Variables:** SLD010H or SLD012 (hours of sleep per night on weekdays/workdays)
- **Cycles:** 2005-2018
- **Definition:** Sleep duration was binned into four right-exclusive categories.
- **Groups:** 1-<5 h, 5-<7 h, 7-<9 h, ≥9 h
5. Sleep Frailty

- **Variables:** All sleep questionnaire variables (SLQ/SLD columns: disorders, problems, behaviors)
- **Cycles:** 2005-2008
- **Computation:** Invalid responses (7, 9, 77, 99) were replaced with median values. Missing binary indicators (e.g., SLQ050, SLQ060, SLQ070A-D) were filled with 0. Sleep duration (SLD010H) was standardized (z-score) and converted to absolute deviation from normal. All sleep variables were scaled (0-1) and averaged to create a composite sleep frailty index.
- **Grouping:** Quartiles were computed, and only the extreme groups were retained.
- **Groups:** Q1 (lowest frailty) vs. Q4 (highest frailty)
6. Alcohol Consumption

- **Variable:** ALQ130 (average drinks per day, past 12 months)
- **Cycles:** 1999-2018
- **Definition:** Consumption was binned into 0-1, 2-4, and >4 drinks/day. Only the extreme groups were analyzed. Invalid responses (77, 99, 777, 999) were excluded.
- **Groups:** 0-1 drink/day vs. >4 drinks/day
7. Social Connections

- **Variables:** SSQ060, SSD061, SSQ061 (number of close friends providing emotional support)
- **Cycles:** 1999-2008
- **Definition:** Binary classification was used based on the presence or absence of close friends. Invalid responses (7, 9, 77, 99) were excluded.
- **Groups:** 0 friends vs. 1+ friends
8. Religious Service Attendance

- **Variable:** SSD044 (times attended religious services in past year)
- **Cycles:** 2005-2008
- **Definition:** Frequency was binned into: Never (0), Sometimes (1-51 times/year), and Weekly (≥52 times/year). Invalid codes (77777, 99999) were excluded.
- **Groups:** Never, Sometimes, Weekly
9. Education Level

- **Variable:** DMDEDUC2 (highest education completed)
- **Cycles:** 1999-2018
- **Definition:** Responses 1-2 were categorized as “No high school diploma,” 3 as “High school graduate/GED,” and 4-5 as “Some college or higher.” Invalid responses (7, 9) were excluded. The middle group (“high school”) was omitted for contrast.
- **Groups:** No high school vs. Some college or higher

For summary statistics on the different groups see Supplementary Table 1.

#### Statistical Analysis

For each exposure group, Kaplan-Meier survival curves were estimated using left-truncated entry (age at enrollment) and right-censored exit (mortality at follow-up). Analyses covered ages 20-110 years. Standard errors for steepness and median lifespan were estimated by bootstrap resampling. Participants were resampled with replacement, survival curves re-estimated, and metrics recalculated. Standard errors correspond to the standard deviation of bootstrap estimates.

### Estimating historic annual births

To approximate the annual number of births occurring in developed countries between 1830 and 1920, we multiplied the world population with the crude birth rate (CBR): *B*(*t*) = *P*(*t*) * *CBR*(*t*)/1000 where *P*(*t*) is the mid-year population and *CBR*(*t*) refers to live births per 1,000 persons. Population data and crude birth rates were taken from *Our World in Data* (OWID)(*68*, *81*). Both population and birth-rate definitions follow WHO conventions for mid-year population and live births per 1,000 persons.

### Historic mortality data

For the historical improvements figure, we used period life table data to construct survival curves for Denmark and Sweden, pooling genders together.

For late-life survival and mortality at extreme ages, we bypassed the smoothed life-table outputs provided by the HMD and instead constructed period life tables directly from the underlying Deaths and Exposure data for each target year. Genders were pooled together. Central death rates were computed as Deaths divided by Exposures. Survival curves were then obtained by standard life-table recursion. To quantify the stochastic uncertainty arising from small sample sizes at ages above 100, we used a Monte Carlo bootstrap with 1000 iterations: in each iteration, death counts were resampled from a Poisson distribution while exposures were held fixed. This yielded 95 percent confidence intervals for the median remaining lifespan.

### Software and computation environment

Analyses were performed in Python 3.11.3, using NumPy 1.26.4, SciPy 1.11.4, pandas 2.3.1, and Matplotlib 3.10.3. Optimization techniques used were SciPy optimize.curve_fit and SciPy optimize.minimize with L-BFGS-B.

## Data Availability

This study does not generate original primary data.

Mortality statistics for Danish, Swedish, French, and Japanese populations are available from the Human Mortality Database(*64*).

NHANES data is available at https://wwwn.cdc.gov/nchs/nhanes/ (NHANES 1999-2000, NHANES 2001-2002, NHANES 2003-2004, NHANES 2005-2006, NHANES 2007-2008, NHANES 2009-2010, NHANES 2011-2012, NHANES 2013-2014, NHANES 2015-2016, NHANES 2017-2018). Linked mortality data for NHANES participants, with follow-up through 31 Dec 2019, are available at https://ftp.cdc.gov/pub/health_statistics/NCHS/datalinkage/linked_mortality/.

HGPS survival data, including risk set and number-of-deaths per time period was digitized from Gordon et al(*73*).

World population data are available at Our World in Data, https://ourworldindata.org/world-population-growth. Crude birth rates are also available at Our World in Data, https://ourworldindata.org/grapher/long-run-birth-rate.

Data on centenarians was gathered from two sources. For the top n ranking centenarians in each calendar year, data was taken from https://www.grg-supercentenarians.org/supercentenarians/(72). Data for the analysis of mean remaining lifespan was taken from the International Longevity Database(*82*, *83*)

## Acknowledgements

We thank Leon Peshkin, Sara Hagg, and all members of the Alon lab for helpful discussions.

## Author Contributions

All authors reviewed the manuscript prior to submission. Conceptualization: BS, UA. Methodology: BS, SF, TL, UA. Formal Analysis: BS Funding acquisition: UA Visualization: BS, SF, TL, UA. Writing: BS, SF, TL, UA. Competing interests The authors declare that they have no competing interests

