## Supplementary Information for "Maximal human lifespan in light of a mechanistic model of aging"

### Supplementary Materials

.

#### Methods

##### Saturating-removal model

We used the saturating-removal (SR) mathematical model(*10*) of aging dynamics, and expanded it here to simulate heterogeneous cohorts. The SR model is a stochastic differential equation that describes the dynamics of damage that is assumed to be causally linked to mortality. Its core assumption is that a dominant form of damage, denoted X(t), drives age-related organismal changes. ·

The SR model describes damage X(t) as a balance of production, removal, and noise

$$\frac{dX}{dt} = \eta t - \beta \frac{X}{X+\kappa} + \sqrt{2\epsilon}\xi .$$

$\eta$, $\beta$, $\kappa$, $\epsilon$ are model parameters, while $\xi$ represents Gaussian white noise. $\eta$ has units of time$^{-2}$, $\beta$ of time$^{-1}$, $\varepsilon$ of time$^{-1}$, $\kappa$ is normalized to 1, and X and Xc are in units of $\kappa$. The model is simulated using an Euler-Maruyama numerical scheme with n=$10^6$ agents, with a reflecting boundary condition at $X = 0$. Death occurs upon crossing a critical threshold $X_C$. Extrinsic mortality $m_{ex}$ is incorporated by adding a death probability of $1 - e^{-m_{ex}dt}$ at each timestep, dt. Population heterogeneity was modeled by sampling model parameters from Gaussian distributions with specified means and standard deviations. Survival and mortality curves are computed from the lifespans generated for the simulated cohort.

#### Steepness of survival curves

We define steepness as the median lifespan divided by the interquartile range of the survival curve: $steepness = \frac{t_{median}}{IQR}$. Other definitions are possible, such as using the inverse coefficient of variation $\mu/\sigma$. However, this latter measure is heavily influenced by the distribution tail. In the exposure groups, too few deaths occurred to estimate the tail reliably, which makes tail-sensitive metrics unstable. Using $\frac{t_{median}}{IQR}$ avoids this problem and provides a robust description of the survival curve's shape.

### Supplementary Text

#### S1. Historical changes in Danish mortality statistics

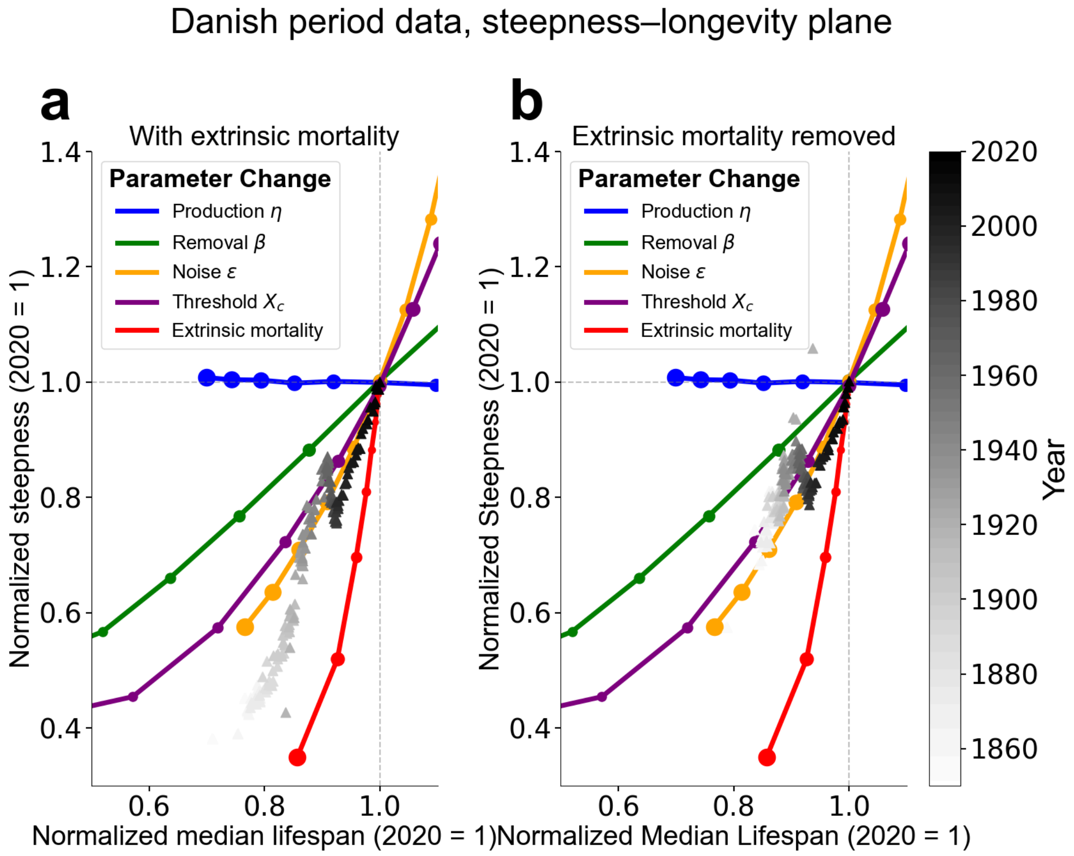


**Supplementary Figure 1. Same as Figure 4 but for Danish period data. a.** Steepness–longevity plot for Danish period survival curves "(males and females pooled), 1840–2000 (shaded by year), normalized to year 2020 data (from HMD). Lines show predicted trajectories for ±70% changes in model parameters. Baseline parameters fitted to Sweden 2019 data: η = 0.49 $year^{-2}$, β = 54.75 $year^{-1}$, κ = 0.5, ε = 51.82 $year^{-1}$, Xc = 17. **b.** Same analysis after removing extrinsic mortality (setting m=0 in the Makeham-Gamma-Gompertz fit).

#### S2. Strehler Mildvan correlation preserves maximum lifespan

Human mortality curves typically obey the Gompertz law of mortality over several decades: $m(t) = ae^{bt}$. The Strehler-Mildvan correlation is the empirical, linear relation between the observed Gompertz slope $b$ and the log of the intercept $log(a)$; $log(a) = B * b + K$ , where $B$ and $K$ are constants. Empirically, the value of the constant B is roughly $B = -105$ years(*28*). For Gompertz mortality, the cumulative mortality is given by $H(t) = \int_0^t m(t)dt = \frac{a}{b}(e^{bt} - 1)$, and the survival function is $S(t) = e^{-H(t)}$. $a$ is of the order $b \approx 0.1\, year^{-1}$, therefore at late ages $t \approx 100\, years$, we may approximate the cumulative mortality as $H(t) = \frac{a}{b}e^{bt}$. Inserting the Strehler-Mildvan equation into the cumulative mortality yields $H(t) = \frac{e^K}{b}e^{b(t+B)}$. Therefore at age $t = -B \approx 105\, years$, the cumulative mortality converges to a constant value across different cohorts, reflecting a maximal lifespan. The value of the slope $-B$ is thus a proxy for the maximal lifespan across different cohorts. This shows that different groups which all respect the same Strehler-Mildvan correlation will have similar maximal lifespan constraints. Thus introducing heterogeneity into parameters which obey the Strehler-Mildvan correlation is not expected to significantly affect maximal lifespan.

#### S3. Maximal lifespan scales in Gompertz hazard as $\ln(\ln N)$

The survival function for Gompertz mortality $m(t) = ae^{bt}$ is given by $S(t) = e^{-H(t)} = \exp(-\frac{a}{b}(e^{bt} - 1))$. For a cohort of size $N$, maximal lifespan is the time for which $S(t_{max}) = \frac{1}{N}$. Taking the same approximation for late ages, $H(t) \approx \frac{a}{b}e^{bt}$, and solving for $t_{max}$ yields

$t_{max} = \frac{1}{b}\ln(\ln N) + \frac{1}{b}\ln\frac{b}{a}$. (1)

A similar scaling occurs in simulations of the SR model, as seen in Figure 5a.

#### S4. Analytical properties of the SR model

##### Derivation of the SR potential

The SR model equation $\frac{dX}{dt} = \eta t - \beta\frac{X}{X+\kappa} + \sqrt{2\epsilon}\xi$ can be interpreted as the over-damped stochastic motion of a particle confined to a potential well:

$\frac{dX}{dt} = -\frac{d}{dx}U(x,t) + \sqrt{2\epsilon}\xi$, where the potential is given by

$$U(x,t) = -\int dx(\eta t - \beta\frac{X}{X+\kappa}) = (\beta - \eta t)X - \beta\kappa\ln(\kappa + X). \qquad (2)$$

#### Quasi-steady-state

While the particles are still confined within the potential well (for $t < \frac{\beta}{\eta} \approx 100\,years$), a quasi-steady-state approximation can be applied to estimate the probability distribution $P(x,t)$. At quasi-steady-state, $X_{st} = \frac{\eta\kappa t}{\beta-\eta t}$. The stationary distribution is given by the Boltzmann distribution $P(x) \propto \exp(-\frac{U(x)}{\epsilon})$. Therefore,

$P(x) \propto \exp(-\frac{(\beta-\eta t)x}{\epsilon})(\kappa + X)^{\frac{\beta\kappa}{\epsilon}}$.

The normalization constant is found by integrating over the allowed range, from 0 to Xc. When age approaches 100, the steady state approximation is no longer valid.

#### Deriving the SR exponential mortality slope

During quasi-steady-state, the SR mortality rate can be estimated using Kramers' approximation for the first-passage time(*84*):

$$m(t) \approx \frac{\sqrt{U''(X_{st})U''(X_C)}}{2\pi}\exp(-\frac{U(X_C)-U(X_{st})}{\epsilon}),$$

where the effective potential was derived above. The curvature around steady-state is $U''(X_{st}) = \frac{(\beta-\eta t)^2}{\kappa\beta}$ while the curvature around $X_c$ is taken as an unknown constant $U''(X_C) = \omega_{Xc}$. This yields the following mortality rate:

$$m(t) \approx \frac{\sqrt{\omega_{Xc}}}{2\pi}(\kappa + X_C)^{-\frac{\kappa\beta}{\epsilon}}(\kappa\beta)^{-\frac{\kappa\beta}{\epsilon}-0.5}(\beta - \eta t)^{-\frac{\kappa\beta}{\epsilon}+1}e^{-\frac{X_C\beta}{\epsilon}}e^{\frac{(\kappa+X_C)\eta t}{\epsilon}}. \qquad (3)$$

Mortality thus rises exponentially with a slope given by $b = \frac{(\kappa+X_C)\eta}{\epsilon} \approx \frac{\eta X_C}{\epsilon}$

#### SR mortality intercept

The mortality intercept (mortality at $t = 0$) for the SR model can be derived exactly. Here, we present the derivation where $\kappa = 0$, as its solution is easy to interpret, requires no cumbersome algebra, and captures the parametric dependence of the model (up to a factor of O(1)). The general case for nonzero κ can be readily obtained using the same derivation steps, although the algebra is messier.

At time $t = 0$, the (fully saturated, k=0) model is $\frac{dX}{dt} = -\beta + \sqrt{2\epsilon}$.

Assuming quasi steady state, $P(x) = Ae^{-\frac{\beta X}{\epsilon}} = \frac{\beta}{\epsilon(1-e^{-\frac{\beta X_C}{\epsilon}})}e^{-\frac{\beta X}{\epsilon}}$, where the constant $A$ is found by normalizing the distribution to 1. For human parameters, $1 \gg e^{-\frac{\beta X_C}{\epsilon}}$, so we approximate $P(x) \approx \frac{\beta}{\epsilon}e^{-\frac{\beta X}{\epsilon}}$.

The initial mortality rate is equivalent to the flux out of $X = X_C$:

$$m(t = 0) = J_{out} = -\epsilon\frac{dP}{dx}|_{X=X_C} = \frac{\beta^2}{\epsilon}e^{-\frac{\beta X_C}{\epsilon}}. \qquad (4)$$

#### Changes in Xc and ϵ preserve the Strehler-Mildvan correlation

Based on the above, the approximate expression for the mortality rate of the SR model is $m_{SR}(t) \sim \frac{\beta^2}{\epsilon} e^{-\frac{\beta X_c}{\epsilon}} e^{\frac{\eta X_c}{\epsilon} t} = A exp(b\, t)$. Indeed we find that $b \propto log(a) \propto X_c, \frac{1}{\epsilon}$, preserving the Strehler-Mildvan correlation. In contrast, changes in damage repair (β) change intercept alone while changes in damage production (η) change slope alone. Both thus break the SM correlation.

The SR model also provides predictions for the empirical parameters B and K of the Strehler-Mildvan correlation $log(a) = B * b + K$:

$$log(m_{SR}(0) * year) = log(\frac{\beta^2}{\epsilon} * year) - \frac{\beta X_c}{\epsilon} = B\frac{\eta X_c}{\epsilon} + K \ . \quad (5)$$

We thus identify $B = -\frac{\beta}{\eta}$ and $K = log(\frac{\beta^2}{\epsilon} * year)$ . The empirical value found for $B \approx 105\, years$ based on mortality curves across cohorts are in agreement with the $\frac{\beta}{\eta} \approx 100\, years$ parameter values based on individual fits to human survival curves.

##### Heterogeneity in damage production (η) and damage removal (β) flatten late-life survival curves

Here we show that large variations in damage production (η) and removal (β) flatten out the late-life survival curves. The general insight is that the mortality rate of those surviving to late age $t$ is dominated by those whose repair rate β is proportional to the total damage production accumulated over time ηt. Large parameter heterogeneity in η leads to a constant mortality rate, while heterogeneity in β leads to an parabolic increase in mortality $\sim t^2$.

With heterogeneous parameters, the total cohort survival curve is given by

$$S_{tot}(t) = \int P(\theta) S(\theta, t) d\theta,$$

where $P(\theta)$ is the parameter distribution for general parameter θ and $S(\theta, t)$ is the survival function at time $t$ for individuals with parameter θ. Assuming Gompertz mortality (and looking at late times):

$$S_{tot}(t) = \int P(\theta)\ \exp(-\frac{a(\theta)}{b(\theta)} e^{b(\theta)t})\, d\theta = \int e^{\Phi(\theta)} d\theta, \quad (6)$$

where $\Phi(\theta) = \ln P(\theta) - \frac{a}{b} e^{bt}$. This integral represents the 'prior' $P(\theta)$, weighted by the survival function. The survival function is essentially 0 when $\frac{a}{b} e^{bt} > 1$, and therefore can be thought of as a step-function which 'kills' all contributions for large values of $\frac{a}{b} e^{bt}$. Therefore, we apply the saddle-point approximation to the integral in order to find the θ which maximally contributes to the integral. When substituting the previous results for the SR Gompertz mortality,

$$\frac{d\Phi}{d\theta} = 0 \rightarrow \frac{d}{d\theta}[\ \ln P(\theta^\star) - \frac{\beta^2}{\eta X_c} e^{-\frac{\beta X_c}{\epsilon}} e^{\frac{\eta X_c}{\epsilon} t}\ ] = 0.$$

Solving for $\theta^{\star}$ gives the point of maximum contribution. The population mortality rate can thus be approximated as $m(t, \theta^{\star})$ - the mortality rate for individuals with parameter $\theta^{\star}$.
Below we separately analyze the results for normal distributions in damage production (η) and damage removal (β).

#### Damage production (η) heterogeneity

We are interested in finding the η value which contributes most to late-life survival. The above expression evaluates to (for large $t \gg \frac{\epsilon}{\eta\, X_C}$):

$$-\frac{\eta-\bar{\eta}}{\sigma^2} - e^{-\frac{\beta X_c}{\epsilon}} e^{\frac{\eta X_c}{\epsilon}t} \frac{\beta^2}{\eta\epsilon}\left(t + \frac{\epsilon}{\eta\, X_C}\right) = 0 \rightarrow -\frac{\eta-\bar{\eta}}{\sigma^2} - e^{-\frac{\beta X_c}{\epsilon}} e^{\frac{\eta X_c}{\epsilon}t} \frac{\beta^2}{\eta\epsilon} t \approx 0$$

Taking the logarithm,

$$\ln\left(-\frac{\eta-\bar{\eta}}{\sigma^2}\right) = -\frac{\beta X_c}{\epsilon} + \frac{\eta X_c}{\epsilon}t + \ln\left(\frac{\beta^2}{\eta\epsilon}t\right) \rightarrow \frac{\eta X_c}{\epsilon}t - \frac{\beta X_c}{\epsilon} = \ln(f(t,\eta))$$

At large $t$, $\frac{\eta X_c}{\epsilon}t - \frac{\beta X_c}{\epsilon}$ must be finite in order for the equality to hold. Therefore, $\eta^{\star} \propto \frac{\beta}{t} + C$. The mean value of $\eta^{\star}$ that contributes to late-life survival scales as $1/t$. Inserting this value into the expression for the mortality rate to find the population mortality at time $t$:

$$m(t) \propto e^{\frac{X_c}{\epsilon}(\beta - \eta^* t)} \approx const$$

In Supplementary Figure 3a-c, we simulate the model with 20% heterogeneity and show that this relation holds.

#### Damage removal (β) heterogeneity

We are interested in finding the β value which contributes most to late-life survival. The above integral evaluates to (for $\frac{\beta X_c}{\epsilon} \gg 1$):

$$-\frac{\beta-\bar{\beta}}{\sigma^2} + e^{-\frac{\beta X_c}{\epsilon}} e^{\frac{\eta X_c}{\epsilon}t} \frac{\beta}{\eta X_c}\left(-2 + \frac{\beta X_c}{\epsilon}\right) = 0 \rightarrow -\frac{\beta-\bar{\beta}}{\sigma^2} + e^{-\frac{\beta X_c}{\epsilon}} e^{\frac{\eta X_c}{\epsilon}t} \frac{\beta^2}{\eta\epsilon} \approx 0$$

Taking the logarithm,

$$\ln\left(\frac{\beta-\bar{\beta}}{\sigma^2}\right) = \ln\left(\frac{\beta^2}{\eta\epsilon}\right) - \frac{\beta X_c}{\epsilon} + \frac{\eta X_c}{\epsilon}t.$$

As before, in order to ensure that the equality holds for large t, we get the same scaling as before: $\beta = \eta t + C$. This again ensures that at late times, $m(t) \propto \frac{\beta^2}{\epsilon} e^{\frac{X_c}{\epsilon}(\beta - \eta^* t)} \propto t^2$. Mortality continues to rise, but as a power-law. In Supplementary Figure 3d-f, we simulate the model with 20% heterogeneity and show that this relation holds.

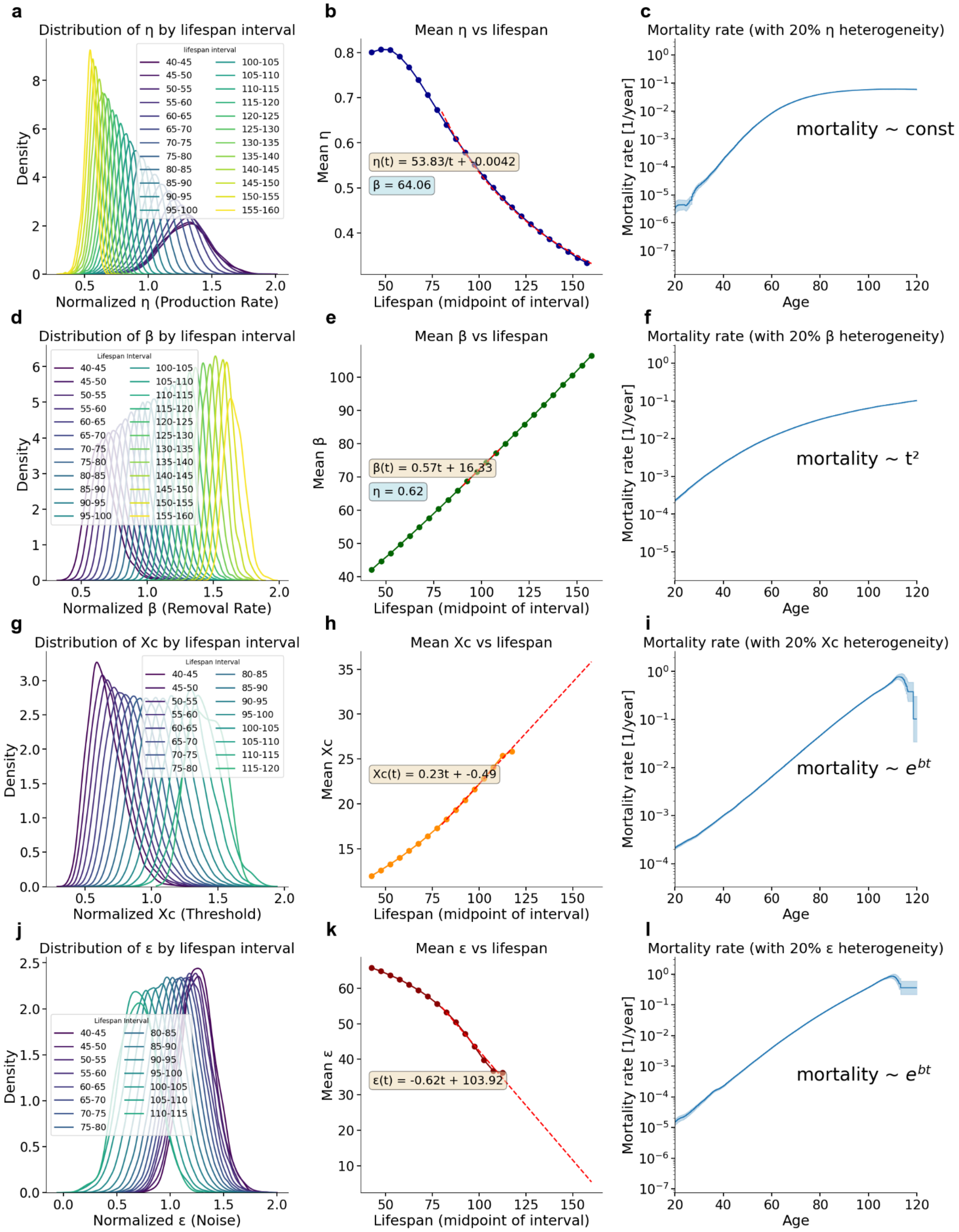


**Supplementary Figure 2. Effect of parameter heterogeneity on late-life mortality in the SR model. Top row:** Analysis of a simulated cohort with 20% variation in the damage production rate (η). **a.** The distribution of η for individuals dying in specific age intervals shifts progressively toward lower values for longer lifespans. **b.** The mean η of survivors scales inversely with lifespan (η(t) proportional to β/t) **c.** This 1/t scaling cancels the time dependence in the Gompertz exponent, producing a hazard rate that approaches a constant (flat) value at late ages. **Second row:**

Analysis of a cohort with 20% variation in the removal rate (β). **d.** The distribution of β shifts toward higher values for longer-lived subpopulations. **e.** The mean β scales linearly with lifespan (β(t) proportional to ηt), acting as a selection force that counterbalances the exponential rise in damage. **f.** The exponential Gompertz hazard becomes algebraic growth (hazard proportional to $t^2$), resulting in pronounced flattening of the mortality curve. **Third row:** Analysis of a cohort with 20% variation in the damage threshold (Xc). **g.** The distribution of Xc shifts toward higher values for longer lifespans. **h.** The mean Xc of survivors increases linearly with lifespan. **i.**The mortality rate maintains exponential growth throughout most of adulthood, with deceleration occurring only at extreme ages. **Fourth row:** Analysis of a cohort with 20% variation in the noise parameter (ε). **j.** The distribution of ε shifts toward lower values for longer-lived subpopulations. **k.** The mean ε decreases linearly with lifespan as individuals with high noise are eliminated early. **l.** Similar to threshold variation, noise heterogeneity generally preserves the exponential Gompertz hazard. Baseline parameter values for all simulations: $\eta = 0.62$ years$^{-2}$, $\beta = 64$ years$^{-1}$, $\kappa = 0.5$, $\varepsilon = 52$ years$^{-1}$, Xc = 18.

#### Derivation of asymptotic mortality rate increase

We are interested in the mortality rate of the model in the asymptotic regime t > β/η. In this regime, stability is already lost because the potential well has flattened, and damage accelerates toward the threshold. For asymptotic times, η ≫ β, which allows us to approximate the model as:

$$\frac{dx}{dt} = \eta t + \sqrt{2\epsilon}\xi$$

The Fokker-Planck equation for this process is:

$$\frac{dP(x,t)}{dt} = -\eta t \frac{dP(x,t)}{dx} + \epsilon \frac{d^2P(x,t)}{dx^2}$$

With $P(x, t)$ being the probability distribution to observe damage $x$ at time $t$.

Without boundary conditions and initial condition of $P(x, 0) = \delta(x)$, the solution is the Green function

$$P(x, t) = \frac{1}{\sqrt{4\pi\epsilon t}} \exp\left[\frac{-(x - \frac{\eta t^2}{2})^2}{4\epsilon t}\right].$$

The full SR model has a reflective boundary condition at $x = 0$. However, since the production term rises with age and is unopposed by removal, most of the probability density at long times is at high values of $x$ and one can thus neglect the boundary condition at $x = 0$. We therefore define the cumulative density up to $x$ as

$$S(x, t) = \int_{-\infty}^{X_c} P(x, t)dx = \frac{1}{2}(1 - Erf(\frac{\eta t^2 - 2X_c}{4\sqrt{\epsilon t}})).$$

Resulting in the asymptotic mortality

$$m(t) = -\frac{d\ln S}{dt} = \frac{3}{16}\left(\frac{\eta^2}{\epsilon}\right)t^2 + \frac{\eta X c}{4\epsilon} - \frac{3}{2t} + O\left(\frac{1}{t^2}\right) \sim \frac{3}{16}\left(\frac{\eta^2}{\epsilon}\right)t^2. \quad (7)$$

Thus, in the absence of heterogeneity, the mortality rate continues to increase algebraically rather than exponentially at late times.

#### S5. Parameter constraints and analytical properties of Fedichev & Gruber's minimal model of aging

Fedichev & Gruber's minimal model of aging(*77*) describes the dynamics of the critical, slow mode of a composite health matrix, described by the following stochastic differential equations:

$$\dot{z}_0 = \beta Z(t) - (\epsilon_0 - \beta' Z(t))z_0 + g z_0^2 + J_0 + \sqrt{2D_0}\xi, \quad (8)$$

$$\dot{Z} = \gamma$$

where $\xi$ is Gaussian white noise. In contrast to the SR model, the removal rate declines linearly with time, with an effective rate of $\epsilon_{eff} = \epsilon_0 - \beta' Z(t)$. Death is modeled as occurring when the mode crosses the protective barrier. The model undergoes a saddle-node bifurcation roughly at time $t_{max} \approx \frac{\epsilon_0}{\beta'\gamma}$, defining the organism's maximal lifespan. The corresponding potential $U$ is given by $U(z_0, t) = -z_0(\beta Z(t) + J_0) + \frac{z_0^2}{2}\epsilon_{eff} - \frac{g z_0^3}{3}$, and initial barrier height is $U_0 = \frac{\epsilon_0^3}{6g^2}$. As in the SR model, mortality can be approximated by Kramers' escape rate: $h(t) \sim e^{-\frac{\Delta U}{D_0}}$, where $\Delta U$ is given by: $\Delta U(t) = \int_{z_{0,min}}^{z_{0,max}} (-\beta Z(t) + \epsilon_{eff} z_0 - g z_0^2) dz_0$. By shifting $z_0$ to be the mid point of the parabola, one can evaluate the integral, yielding the exact relation for the drop in barrier height,

$$\Delta U(t) = \frac{\epsilon_{eff}^3}{6g^2}\left(1 - \frac{4g\beta\gamma t}{\epsilon_{eff}^2}\right)^{3/2}. \quad (9)$$

For $\frac{4g\beta\gamma t}{\epsilon_{eff}^2} \ll 1$, the potential barrier can be approximated as $\Delta U(Z) \approx \frac{\epsilon_{eff}^3}{6g^3}\left(1 - 6\frac{g\beta Z}{\epsilon_{eff}^2}\right)$. Keeping only terms linear in $Z$ (approximating for small damage erosion), we find $\Delta U(Z) \approx \frac{\epsilon_0^3}{6g^2} - Z\left(\frac{\epsilon_0\beta}{g} + \frac{\epsilon_0^2\beta'}{2g^2}\right)$. Therefore, the approximate mortality rate is given by

$$m(t) \approx e^{-\frac{\epsilon_0^3}{6g^2 D_0}} e^{\gamma t\left(\frac{\epsilon_0\beta}{g} + \frac{\epsilon_0^2\beta'}{2g^2}\right)}. \quad (10)$$

The exponential increase in mortality is due to the combined effect of the accumulation of irremovable damage $Z$ and the erosion of removal $\epsilon_{eff} = \epsilon_0 - \beta' Z(t)$.

Thus, changes in β, β' , or γ affect the mortality slope alone, similar to varying the damage production rate (η) in the SR model, and thus do not respect maximal lifespan constraints. Changes in the initial resilience parameter $\epsilon_0$ or the non-linear coupling term $g$ also do not preserve maximal lifespan constraints due to different scaling in the intercept and slope. Conversely, only changes in $D_0$ (noise amplitude) preserve the Strehler-Mildvan correlation and therefore respect maximal lifespan constraints. $D_0$ plays an identical role to noise amplitude ϵ in the SR model.

#### Late-life mortality slow-down

Recall the expression for $\Delta U(t) = \frac{1}{6g^2}(\epsilon_{eff}^2 - 4g\beta\gamma t)^{3/2}$.
To understand late-life mortality behavior, we examine the two extreme cases:

a) Where damage accumulation is negligible ($\beta \approx 0$)
b) Where removal rate decline is negligible ($\beta' \approx 0$)

a) $\Delta U(t) = \frac{1}{6g^2}\epsilon_{eff}^3 \propto (1 - \frac{t}{t_{max}})^3$. The Gompertz slope is equal to the change in barrier height, and is proportional to $|\frac{d\Delta U}{dt}| \propto (1 - \frac{t}{t_{max}})^2$. As $t \to t_{max}$, the slope thus drops off quickly. b) $\Delta U(t) = \frac{1}{6g^2}(\epsilon_0^2 - 4g\beta\gamma t)^{3/2} \approx (1 - \frac{t}{t_{max}})^{3/2}$. The Gompertz slope is equal to the change in barrier height, and is proportional to $|\frac{d\Delta U}{dt}| \propto (1 - \frac{t}{t_{max}})^{1/2}$. The slow-down in this case is weaker than in the previous case.
The relative strengths of β and β' therefore impact the slow-down behavior leading up to $t_{max}$.

#### Coefficient of variation of $z_0(t)$

The mean of $z_0$ behaves as $< z_0 > \approx \frac{\beta Z + J_0}{\epsilon_{eff}}$, and the standard deviation as $\sqrt{var(z_0)} \approx \sqrt{\frac{D_0}{\epsilon_{eff}}}$.
Therefore, the coefficient of variation behaves as $CV = \frac{\sigma}{<z_0>} = \frac{\sqrt{D_0\epsilon_0}}{\beta\gamma t + J_0}\sqrt{1 - t/t_{max}}$. This expression can be divided into two regimes:

1. $t \ll t_{max}$: during this phase, CV rises decreases hyperbolically with time $CV \sim \frac{1}{t}$
2. $t \sim t_{max}$: during this phase, CV approaches 0 as $CV \sim \sqrt{t_{max} - t}$

CV thus decreases with time.

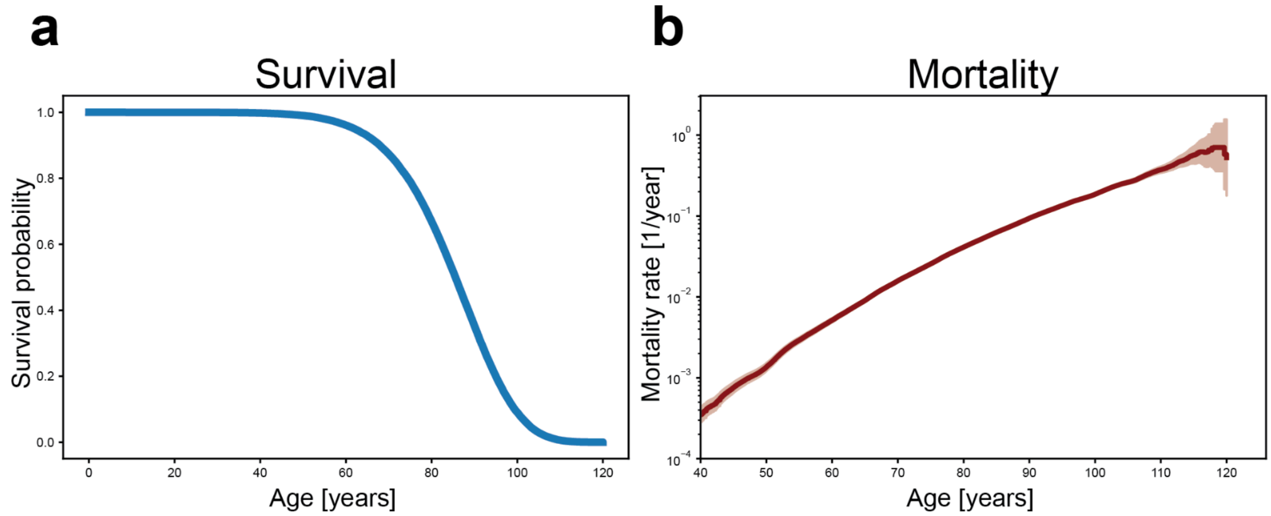


**Supplementary Figure 3. Fedichev & Gruber's minimal aging model. a.** Survival probability for a representative parameter set. The chosen values reproduce the characteristic shape of human survival curves. **b.** Mortality rate for the same parameter set. The model yields the expected Gompertz-like exponential increase. Baseline parameters: $\beta' = 0.013$, $\varepsilon_0 = 4$, $\gamma = 1$, $\beta = 0.015$, $g = 0.8$, $D_0 = 1.1$.

#### S6. NHANES exposure groups summary statistics

| Topic | Group | N | Deaths observed | Median lifespan (years) | Steepness |
|---|---|---|---|---|---|
| Alcohol | 0-1 drink/day | 11241 | 1738 | 86 | 6.1 |
| Alcohol | >4 drinks/day | 4963 | 430 | 78 | 3.9 |
| Church … | never | 2415 | 817 | 80 | 4.7 |
| Church … | sometimes | 1882 | 438 | 83 | 4.9 |
| Church … | weekly | 1890 | 582 | 85 | 5.7 |
| Diet | Good | 12236 | 1656 | 84 | 5.6 |
| Diet | Poor | 27271 | 2586 | 83 | 4.9 |
| Educati… | no highschool | 15116 | 3710 | 80 | 4.0 |
| Educati… | some college | 27001 | 3163 | 84 | 5.2 |

| | | | | | |
|---|---|---|---|---|---|
| Income | Q1 (Lowest) | 7897 | 763 | 78 | 3.5 |
| Income | Q2 | 7770 | 993 | 79 | 4.2 |
| Income | Q3 | 7783 | 788 | 84 | 5.6 |
| Income | Q4 (Highest) | 7784 | 486 | 86 | 7.8 |
| Number… | 0 friends | 637 | 288 | 78 | 4.1 |
| Number… | 1+ friends | 13031 | 5571 | 83 | 4.9 |
| Physical… | No Activity | 12510 | 2147 | 80 | 4.4 |
| Physical… | Some Activity | 23951 | 1356 | 86 | 5.4 |
| Sleep D… | 1-<5 hours | 2092 | 307 | 79 | 3.6 |
| Sleep D… | 5-<7 hours | 12255 | 1251 | 83 | 4.6 |
| Sleep D… | 7-<9 hours | 22189 | 2178 | 84 | 5.6 |
| Sleep D… | ≥9 hours | 5389 | 781 | 78 | 3.5 |
| Sleep Fr… | Q1 (lowest) | 2951 | 614 | 83 | 5.2 |
| Sleep Fr… | Q4 (highest) | 2945 | 532 | 81 | 3.9 |

**Supp. Table 1. Summary statistics for NHANES exposure groups.** *N* is the number of individuals in each exposure group, which varies across groups due to differences in question availability across NHANES survey cycles. "Deaths observed" indicates the number of deaths recorded from the date of questionnaire administration through the end of follow-up (31 Dec 2019). Median lifespan and steepness values were derived from Kaplan-Meier survival curves estimated for each cohort.

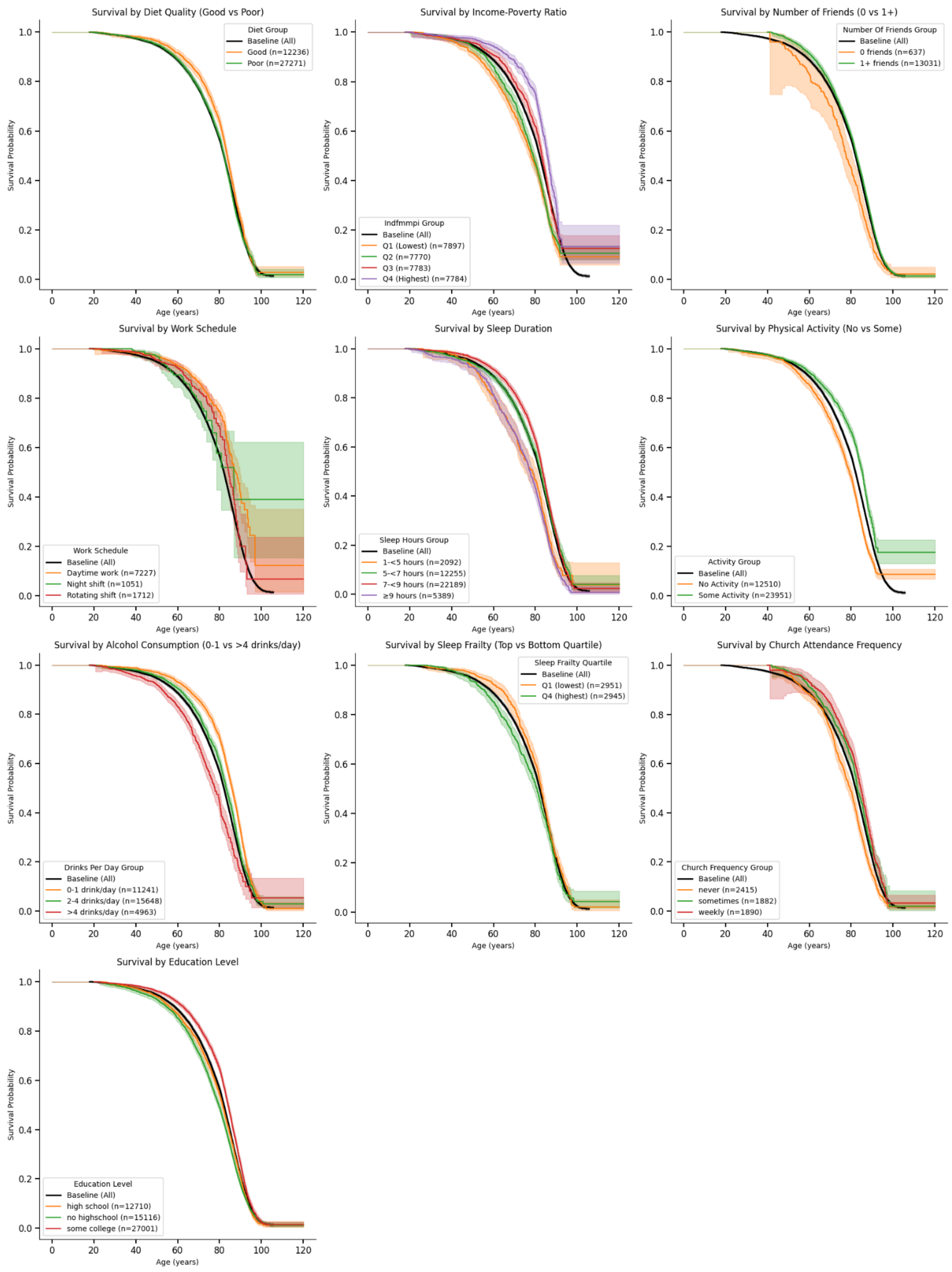


**Supplementary Figure 4. Raw Kaplan-Meier survival curves of the different NHANES exposure groups.** Survival curves displayed here are not corrected for extrinsic mortality.

#### S7. Rank statistics of oldest living individuals

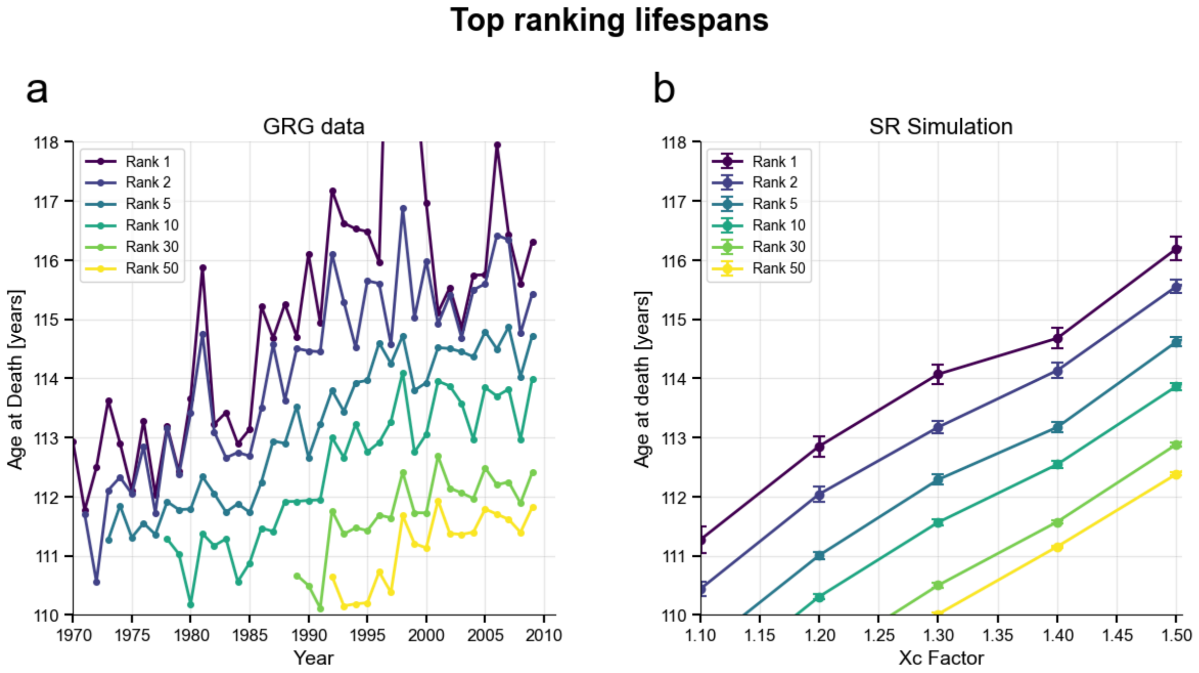


**Supplementary Figure 5. Top ranking lifespans. a.** Top-ranked ages at death from the GRG database **b.** Top-ranked ages at death obtained from the go with the winners SR simulation for different threshold Xc values. Error bars represent the standard error of the mean estimated by bootstrapping. Baseline parameter values: Baseline parameters: $\eta = 0.66$ year$^{-2}$ , $\beta = 62.05$ year$^{-1}$, $\kappa = 0.5$, $\varepsilon = 51.83$ day$^{-1}$, $Xc = 14$, with 20% person-to-person normal variation in threshold Xc.
